# Estimating error rates for single molecule protein sequencing experiments

**DOI:** 10.1101/2023.07.18.549591

**Authors:** Matthew Beauregard Smith, Kent VanderVelden, Thomas Blom, Heather D. Stout, James H. Mapes, Tucker M. Folsom, Christopher Martin, Angela M. Bardo, Edward M. Marcotte

## Abstract

The practical application of new single molecule protein sequencing (SMPS) technologies requires accurate estimates of their associated sequencing error rates. Here, we describe the development and application of two distinct parameter estimation methods for analyzing SMPS reads produced by fluorosequencing. A Hidden Markov Model (HMM) based approach, extends *whatprot*, where we previously used HMMs for SMPS peptide-read matching. This extension offers a principled approach for estimating key parameters for fluorosequencing experiments, including missed amino acid cleavages, dye loss, and peptide detachment. Specifically, we adapted the Baum-Welch algorithm, a standard technique to estimate transition probabilities for an HMM using expectation maximization, but modified here to estimate a small number of parameter values directly rather than estimating every transition probability independently, which should help prevent overfitting. We demonstrate a high degree of accuracy on simulated data, but on experimental datasets, we observed that the model needed to be augmented with an additional error type, N-terminal blocking. This, in combination with data pre-processing, results in reasonable parameterizations of experimental datasets that agree with controlled experimental perturbations. A second independent implementation using a hybrid of DIRECT and Powell’s method to reduce the root mean squared error (RMSE) between simulations and the real dataset was also developed. We compare these methods on both simulated and real data, finding that our Baum-Welch based approach outperforms DIRECT and Powell’s method by most, but not all, criteria. Although some discrepancies between the results exist, we also find that both approaches provide similar error rate estimates from experimental single molecule fluorosequencing datasets.

## Introduction

Proteins are key components of all living organisms, but their roles and functions, and in particular their quantities, processing, and modifications, cannot be fully determined from DNA and RNA sequencing alone. Advancements in protein identification and quantification are much sought after by researchers in the field of single molecule protein sequencing. SMPS technologies apply concepts from DNA and RNA sequencing to protein analysis—including imaging or nanopore-based assays, high parallelism, and single molecule detection—with the hope of achieving the high throughput and sensitivity offered by such approaches [1–5].

Notably, early generations of single molecule DNA sequencing technologies were marked by high error rates, but subsequent optimization allowed these error rates to be driven down. For example, the first commercially-released MinION R7 nanopore sequencers in 2014 from Oxford Nanopore Technologies generally showed error rates at or above 30% error (per nucleotide, per molecule) [6–9], but improvements in chemistry and software rapidly brought error rates down to approx. 15% within just a year or two [10], and continued optimization has reduced the error rates to their current levels of < 5% (e.g. [11,12]). One might reasonably expect a similar trajectory for SMPS technologies, highlighting the importance of accurately estimating the sequencing parameters and error rates in order to facilitate practical applications and ongoing optimization of the methods.

In the SMPS technique known as fluorosequencing (**Figure 1**) [13,14], a biological sample of peptides is acquired, such as from the proteolytic digestion of proteins. The peptides are then covalently labeled with fluorescent dyes on specific types (or groups) of amino acids. The peptides are then immobilized by their carboxy-termini in a microscope flow cell and imaged by total internal reflection fluorescence (TIRF) microscopy. By alternating microscopy with performing Edman degradation chemistry [15], which removes a single amino-terminal (N-terminal) amino acid from each peptide on each sequencing cycle, a time series of images is obtained capturing sequence information for many peptides in a highly parallel and scalable fashion. Signal processing of the resulting images is performed to identify peaks, corresponding to peptides, and quantify their fluorescence across the time series, thus providing the primary raw sequencing reads, each comprising a time series of fluorescence intensities for a single peptide molecule across the course of the sequencing experiment. As these reads capture the sequence positions of the fluorescently labeled amino acids within each peptide, they can then be matched to a reference database in order to identify the most likely peptides, and thus proteins, in the sample. We previously described the algorithm *whatprot* to perform this peptide-read matching process to classify the reads, assuming the set of fluorosequencing parameters for the experiment [16].

**Figure 1.**
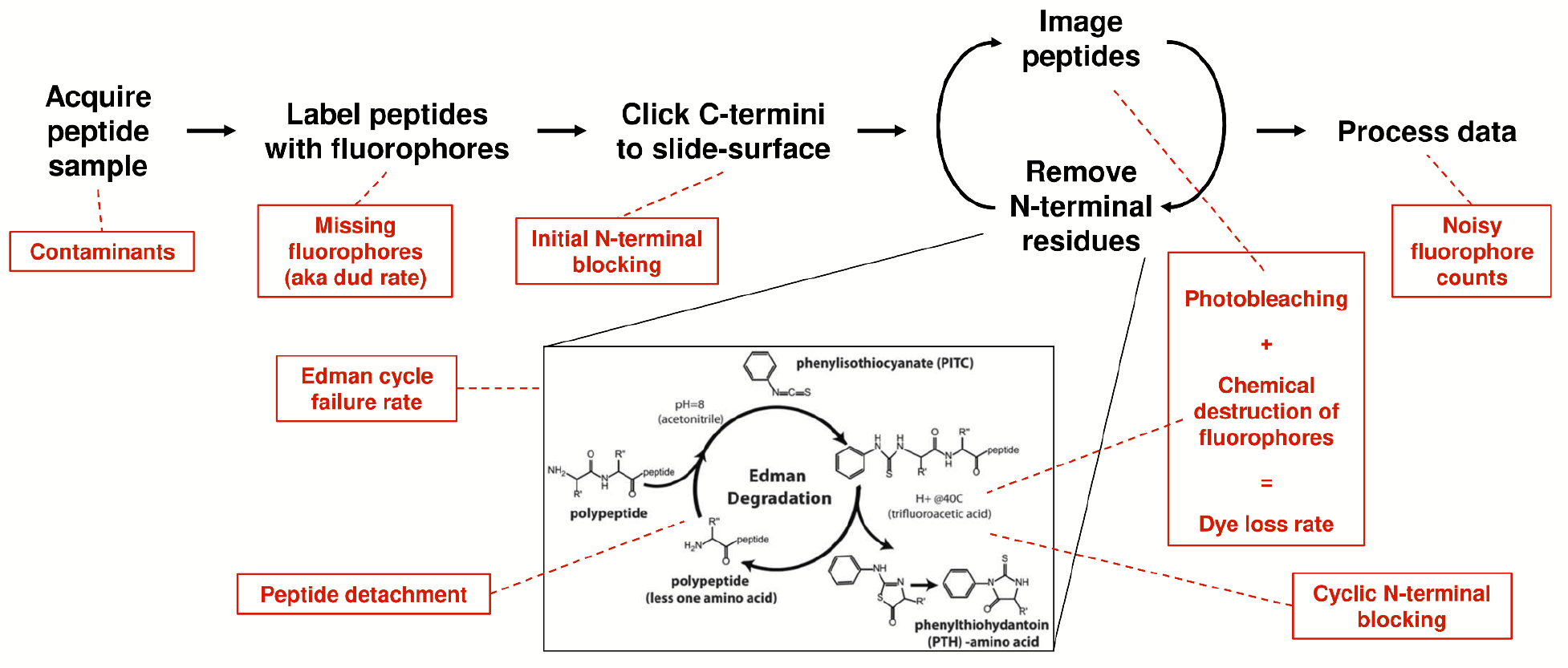
Overview of single molecule protein fluorosequencing with various potential sources of errors highlighted in red. A sample of peptides is acquired, which due to biological factors or collection practices may contain contaminants. These peptides are then labeled, but this process is not expected to be 100% efficient, and a missing fluorophore rate must therefore be considered. Labeled peptides are clicked to the slide surface prior to sequencing; N-terminal modifications and non-specific attachment might potentially occur during this process. The peptides are then imaged and a single amino acid removed from each of their N-termini by Edman degradation, and these two processes iterated to fully sequence the peptides. Imaging photobleaches dyes, while incubation in trifluoroacetic acid (TFA) and phenyl isothiocyanate (PITC)/pyridine can result in chemical destruction of fluorophores. As photobleaching happens at a negligible rate in the imaging conditions used [13] and the dye loss rate is dominated by the contribution of chemical destruction, we combine these effects into a single dye loss rate. Edman degradation is not 100% efficient, and we model failures with the Edman cycle failure rate. We model the potential for blocking peptide N-termini during the course of sequencing, which we call cyclic N-terminal blocking, and also model the detachment of intact peptides (e.g. by non-specific cleavage or washing off of non-specifically attached peptides). Finally, it can be difficult to entirely denoise the precise fluorophore counts due to overlapping intensity distributions, so we additionally recognize an error contribution from mis-assigned fluorophore counts.

Whatprot models the sequencing of each peptide by a distinct HMM, an example of which is illustrated in **Figure 2**. These HMMs capture separate states for each possible condition of the peptide during sequencing, including the number of successful Edman cycles and the fluorophores remaining. In general, HMMs consist of a set of states and a set of probabilistic transitions between those states. A straightforward way to model fluorosequencing with an HMM is to have a state for every possible condition of the peptide, including the number of remaining amino acids (amino acids not yet removed by Edman degradation), and the combination of fluorophores still present on the peptide. Because each fluorophore can either be present and functioning or not, the number of states in the model grows exponentially with respect to the maximum possible number of fluorophores for the peptide given the experimental conditions.

**Figure 2.**
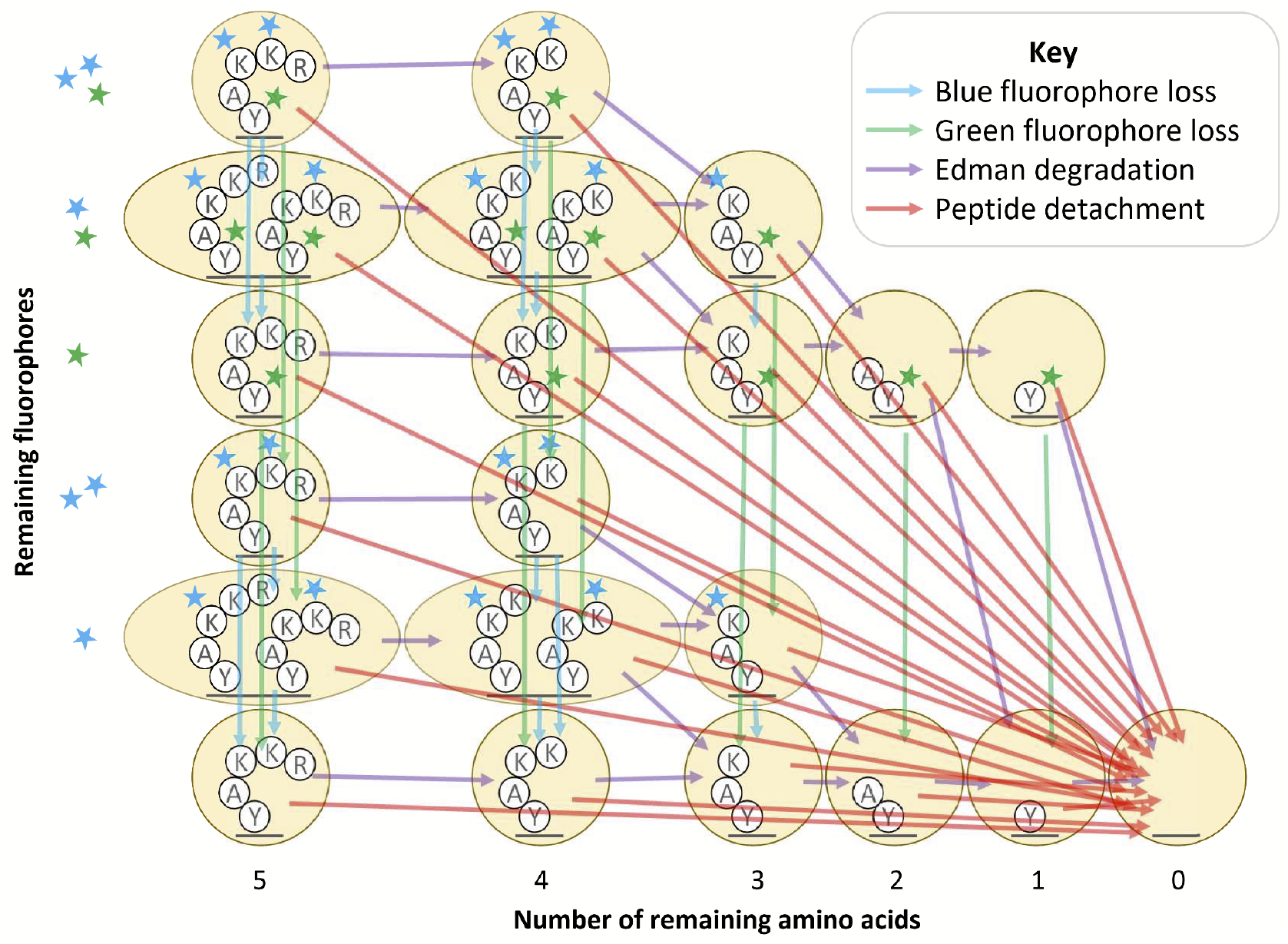
Diagram of factored hidden Markov model for fluorosequencing of an example peptide with three labeled amino acids. This figure is adapted from Figure 6 in [16]. Arrows of a particular color indicate the non-zero entries in a factor of the transition matrix for a particular form of error (see key).

We previously found that significant optimizations of the HMM were necessary to make the forward algorithm tractable to run in practice. In particular, we reduced the state space of the model by merging all states with the same number of successful Edman cycles and fluorophore counts of each color (**Figure 2**, oval states), and we also introduced a factorization of the transition matrix which, as a side effect, isolated the effects of each form of error from the others (**Figure 3**, as discussed below) [16].

**Figure 3.**
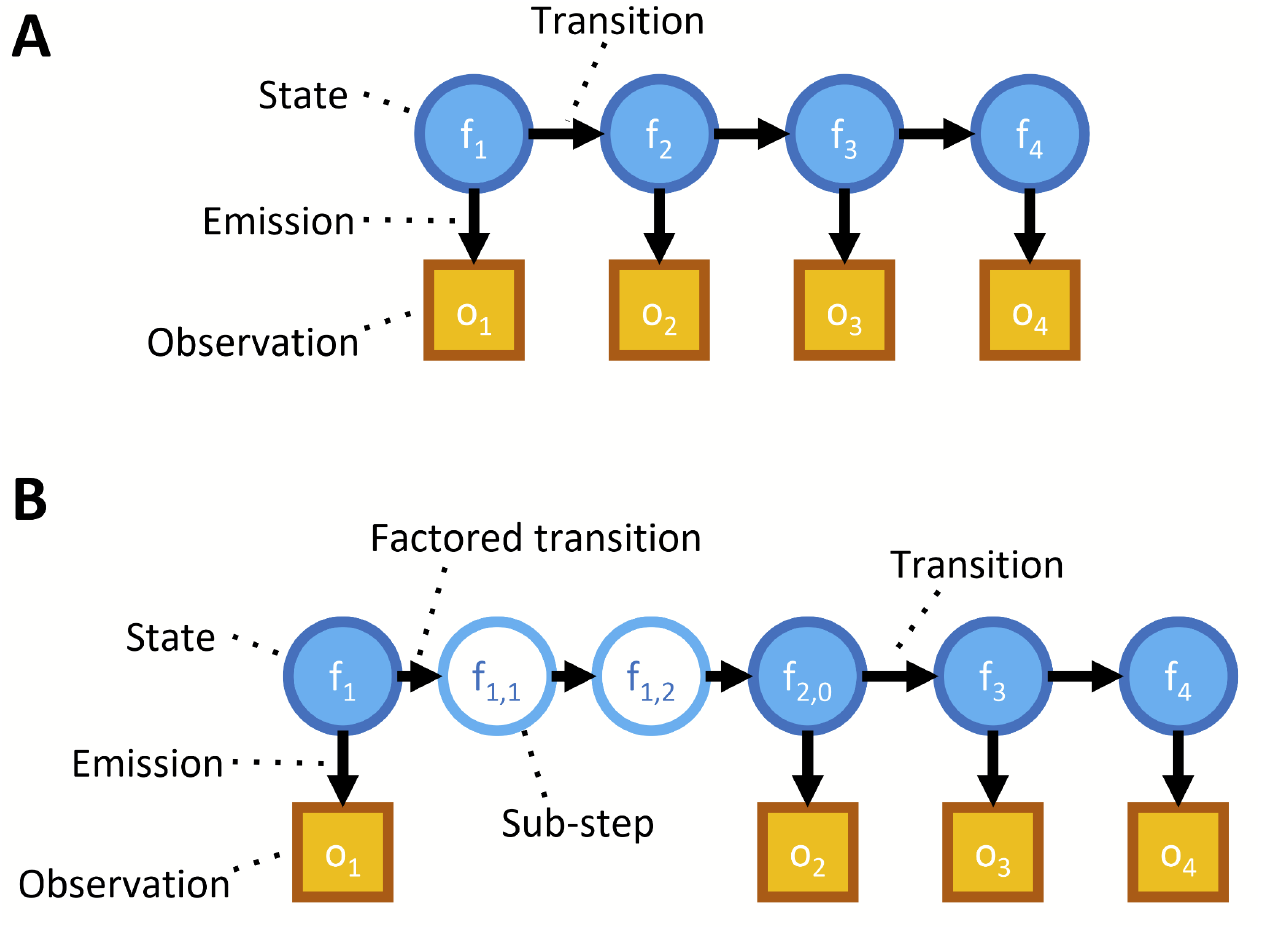
Illustration of HMM factorization. (A) Diagram of a non-factored HMM model. Arrows represent a conditional probability relationship. The transitions between states determine how a state at one time step is probabilistically related to the state at the preceding time step. Emissions represent how the observable data is probabilistically determined by the associated state. (B) Diagram including a factored transition. Breaking a transition into a factored product of sub-transitions introduces “sub-steps”; though not accurate models of any physical states of an actual peptide, these sub-steps prove useful for algorithmic purposes.

Separating forms of error proved fortuitous for parameter fitting, as it allows isolation of each error type. Further, we found that we could perform weighted maximum likelihood estimation, in a modification of the classical Baum-Welch algorithm [17], to estimate sequencing parameters directly. This approach offers some advantages over the classical Baum-Welch algorithm, including greater interpretability and direct ties to the underlying model of the fluorosequencing process, and seems likely to be less prone to overfit the data.

We implemented these new capabilities in parameter estimation as an extension to *whatprot*. We demonstrate that the approach provides accurate estimates of the parameters used for simulated data, correctly identifies experimentally manipulated parameters in actual sequencing experiments, and gives similar parameter estimates to simpler, more general purpose, parameter estimation techniques that are by their nature less precise.

## Methods

### Ordinary Baum-Welch

We estimated parameters for fluorosequencing experiments using a modification to the classical Baum-Welch algorithm. First, we briefly review Baum-Welch [17], a standard technique to determine transition probabilities in an HMM. Dynamic programming is used to determine the probability, given a sequence of observed data, of being in each state at each time step. The probability of each particular transition occurring between each neighboring pair of time steps is also determined. Together, these probabilities allow a weighted maximum likelihood estimate (MLE) to be computed for every transition probability in the model.

Similarly, an estimate for the emission distribution can also be determined. This is done by simply computing a weighted MLE of the hypothetical distribution—in this case a normal distribution—in order to parameterize it, independently for each possible state.

We note that weighting this estimate is necessary to properly account for the fact that a state at one time step may be more likely, and therefore a better source of data, than that same step at a different time step; the more likely time step should have a greater impact on the final result. Additionally, in our model of a peptide undergoing fluorosequencing, many transitions are impossible; these transitions are omitted from the Baum-Welch procedure to improve runtime, as we know their probability to be zero.

Baum-Welch is a form of Expectation Maximization (EM) algorithm, and uses an existing estimate for modeling parameters to compute a better estimate of the modeling parameters. We therefore iterate this process until convergence is achieved. In **Appendix A1**, we provide a full mathematical description of the Baum-Welch algorithm. Additionally, Baum-Welch can be easily generalized to multiple independent and identically distributed sequences of output data, a simple and common modification which we also describe in the appendix.

### Baum-Welch with isolation of error types

In [16] we improved the algorithmic complexity of the forward algorithm by factoring the transition matrix into a product of sparse matrices (**Figure 3**). We also showed that this factorization involved no loss in the accuracy of the model. For parameter estimation, only one peptide needs to be considered, so runtime considerations are of less significance. However, we find that this factorization serves a second purpose: isolation of our forms of error.

The Baum-Welch algorithm must be modified to take advantage of the factored transition matrix. In particular, we consider every step as an indexed series of sub-steps, each pertaining to a particular form of error; further, we can consider each transition as a series of sub-transitions, each pertaining to an application of one of our factors of the original transition matrix. We can then determine the probability of being in each of the states of the model at every sub-step of every time step, as well as the probability of any sub-transition between adjacent sub-steps. We then proceed as in the classical Baum-Welch with these probabilities to estimate the sub-transition probabilities using weighted MLEs.

A more rigorous mathematical description of this modification to the classical Baum-Welch algorithm is provided in **Appendix A2**. This algorithm was never implemented, and is instead considered as a theoretical stepping stone to direct parameter estimation, described in the following section.

### Weighted MLE for parameter estimation

While isolation of error types through factorization of the transition matrix should lead to a better model of fluorosequencing, it falls short of direct estimation of the model parameters. For this reason we made a further modification to Baum-Welch. Instead of estimating each transition probability independently through weighted MLE in every iteration of Expectation Maximization, we use weighted MLE to directly estimate the parameters.

For example, consider the sub-transitions of dye loss for a particular color of fluorophore. If these sub-transitions are considered independently of each other, then their new probability for the next iteration of Expectation Maximization is given by the probability of the sub-transition, which is then normalized by dividing by the probability of being in the starting state at the sub-time step for the sub-transition. We instead consider each probabilistic transition as evidence for the value of an underlying parameter, in this case the per-cycle dye loss rate of the fluorophore.

Weighting of the MLE is necessary because *n*, the number of trials, and *x*, the number of successes (which for this application is counter-intuitively the probability of fluorophore loss, which is in a way a sort of failure), can only be determined probabilistically. However,

Baum-Welch provides the weights for these probabilities. We can take *n* to be a sum of the number of fluorophores for every state at the sub-time step before the sub-transition, weighted by the probabilities of being in each of those states. Similarly, we take *x* to be a sum of the number of fluorophores of each destination state of a sub-transition, weighted by the probabilities of those sub-transitions.

This approach can be generalized to other parameters under consideration. This is done by mapping each starting state and sub-transition to a value which is then fed into a weighted estimate. In **Appendix A3**, we provide a generalized formula for the MLE step of the Baum-Welch algorithm, along with descriptions of the value-mapping procedures used for each type of sub-transition.

### Bias correction for the missing fluorophore rate

The missing fluorophore rates (aka “dud rates”, for brevity), which are the rates of non-fluorescing or absent fluorophores of each color before sequencing starts, have additional challenges. If all fluorophores on a peptide are missing, the peptide will not show up as a read in the sequencing results. This introduces a problematic bias, and if not accounted for in an MLE, the missing fluorophore rates may be estimated significantly lower than they should be (**Figure 4**). We derived an equation for the MLE of a binomial distribution with all instances of all fluorophores missing. Our MLE equation allows no closed-form solution that we know of; however, it does suggest an iterative method (**Appendix A4**).

**Figure 4.**
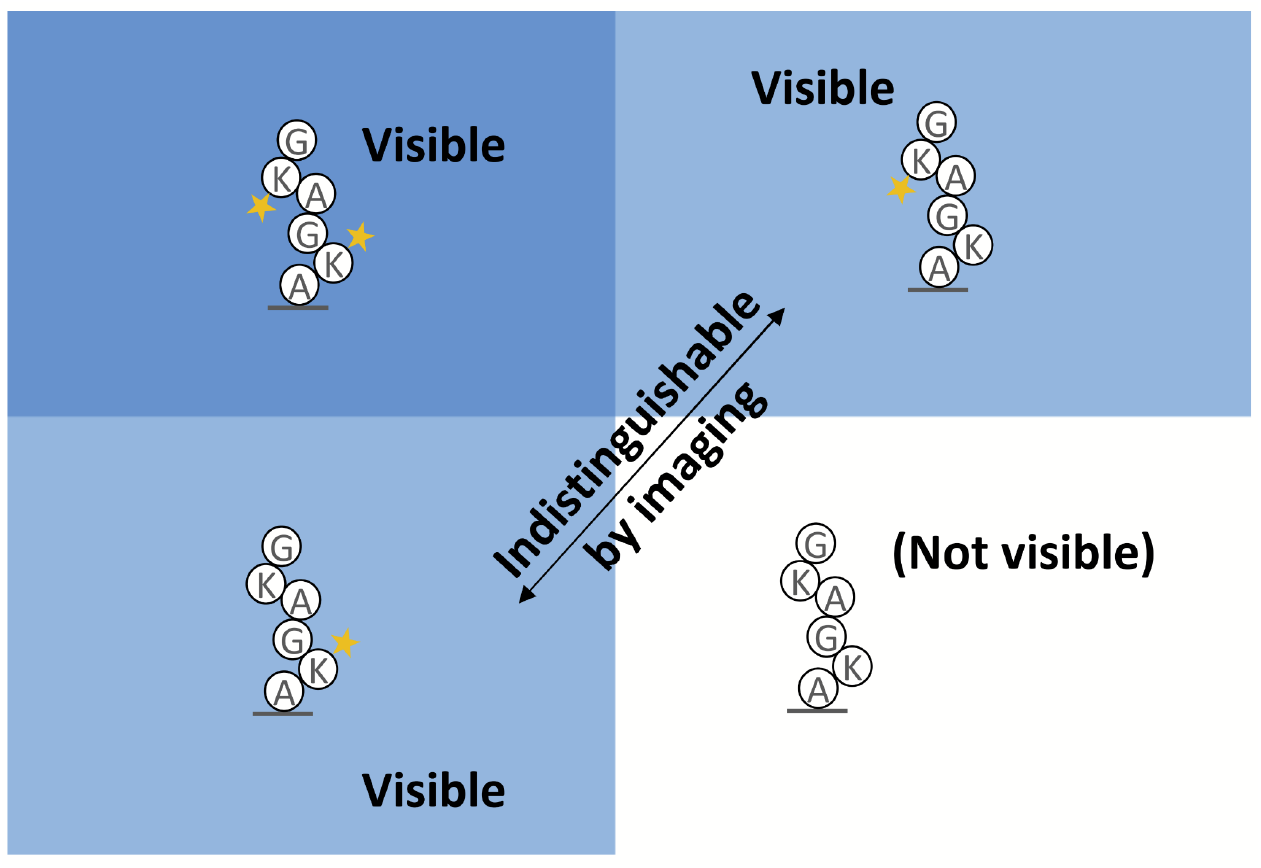
Data is lost due to missing fluorophores. At least one functioning fluorophore must be present when sequencing starts to observe a peptide. This makes naive calculations of the missing fluorophore rate biased, and a correction is needed.

We interleave this iterative bias-correcting process with our implementation of the Baum-Welch algorithm. In particular, the missing fluorophore rates of the previous iteration determine the probability, *x*, that the peptide is missing from the dataset. We then compute *N(x/(1-x))* where *N* is the number of reads. This serves as an estimate of the number of missing reads, which we use for each missing fluorophore rate estimator as an instance of a peptide for which all of its dyes are missing.

### N-terminal blocking

In exploratory work for this publication, we found that parameter estimation results on real datasets required that we consider an additional form of error that was not accounted for in our prior models of fluorosequencing: the N-terminus could have a chemical modification that blocks sequencing, which is either acquired before sequencing starts (initial N-terminal blocking) or while sequencing takes place (cyclic N-terminal blocking). To account for initial and cyclic N-terminal blocking, we adjusted our model, both for purposes of classification (as in [16]) and for parameter fitting as follows:

We first doubled the number of states, in order that each of the former states now maps to two states, one with and one without N-terminal blocking. Edman degradation sub-transitions only apply to the states without N-terminal blocking and have no effect on the states with blocked N-termini. Other sub-transitions and the emission calculations are applied equivalently but separately to both sets of states as before. However, we require a new sub-transition factor of our transition computations to account for N-terminal blocking: with a parameterized probability, a state transition is made from a state to its N-terminally blocked duplicate. This new transition has two variants, one to account for initial N-terminal blocking, and another to account for per-cycle N-terminal blocking (**Figure 5**).

**Figure 5.**
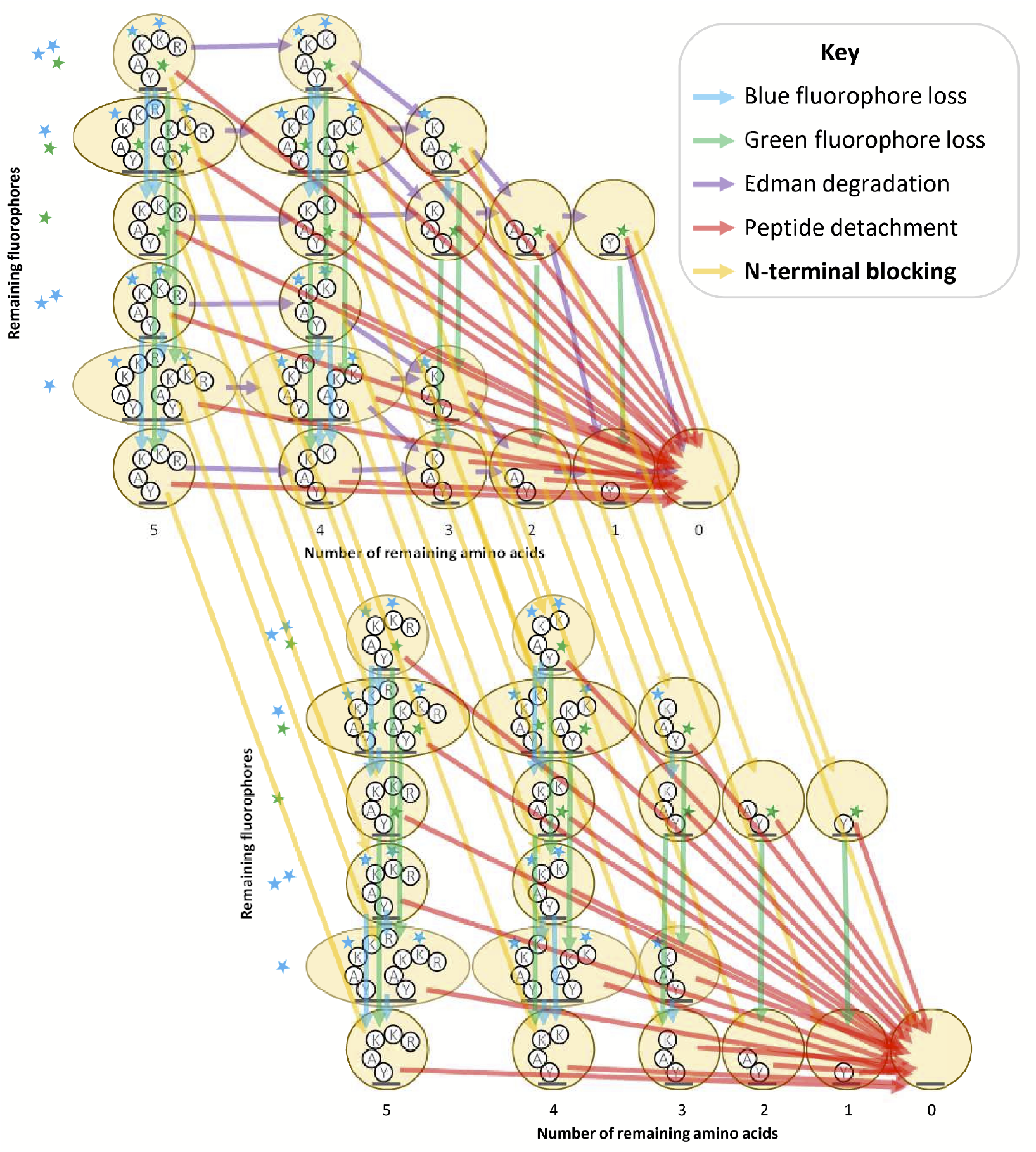
Factored transition matrix diagram including N-terminal blocking. In this diagram we adapted the illustration from **Figure 2** to include N-terminal blocking. In addition to the unblocked states (top-left) described in [16], we found we must also consider blocked states (bottom-right) in which Edman degradation is not possible. An additional transition matrix factor representing N-terminal blocking is needed to describe this behavior (yellow arrows).

### Contaminants and abnormal intensity distributions

While the above approach works well on simulated data, we observed two additional factors affecting actual experimental data: (1) contaminants in the sample that behave like fluorophores, and (2) fluorophore intensity distributions not precisely fitting a normal distribution (see, for example, the histograms in **Figure 6A**).

**Figure 6.**
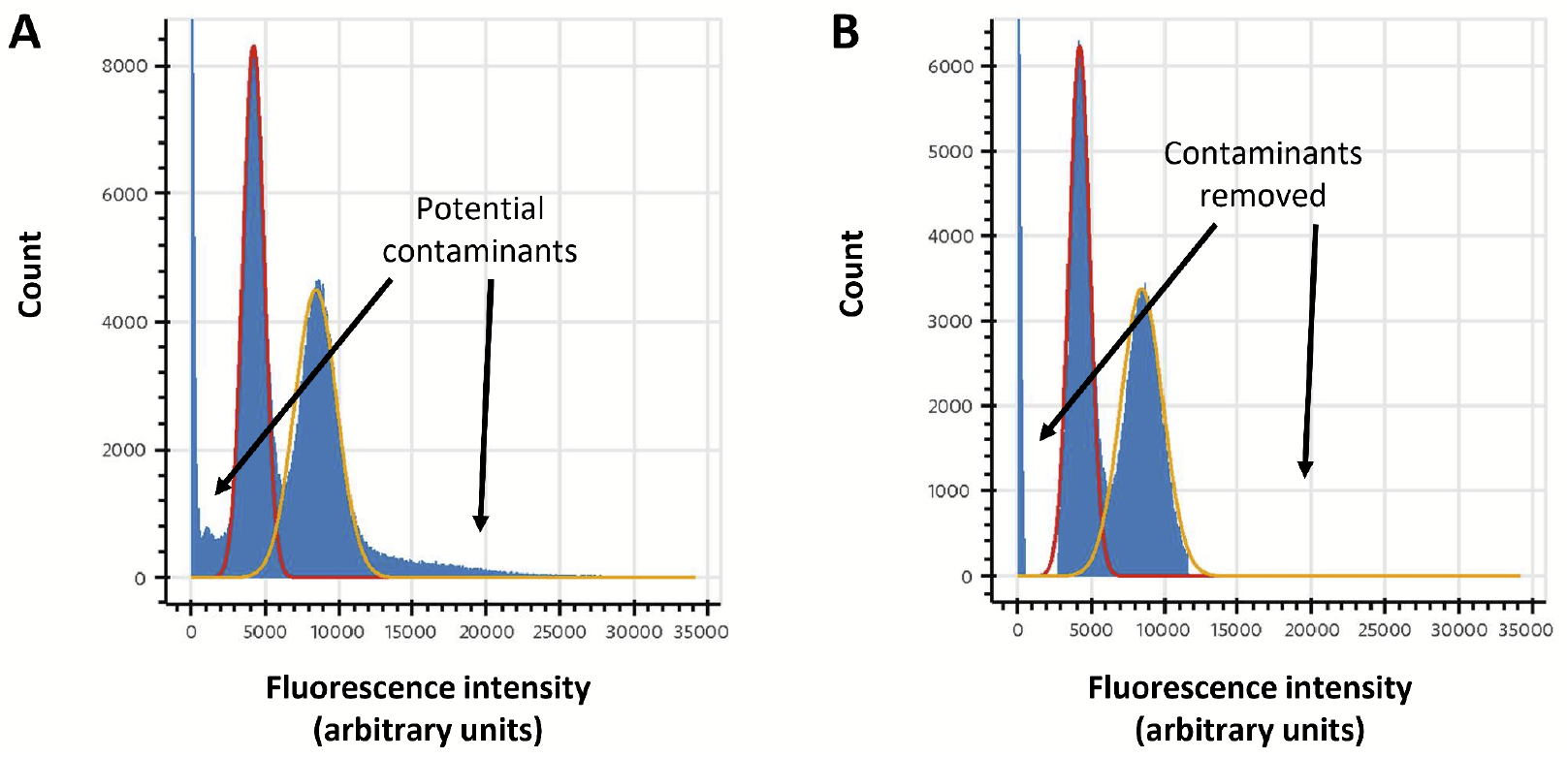
Determination of fluorescence intensity distribution parameters and filtering of likely contaminants. (A) unaltered histogram of intensity values for a NH2-G{azK}*AG{azK}*| peptide sequencing experiment with superimposed normal distributions (red, fit to peak max (μ) and half-width; yellow, expectation from 2*μ). We typically observe some deviation from a normal distribution that can cause challenges with fitting the distribution, often solved in practice either by fitting the max and peak half-width (as in the red curve) or by trial-and-error using expert judgment. (B) clipped data for the same experiment, removing ranges of intensity values likely to be caused by contaminants or signal bleed over from adjacent peaks. Typically, reads are removed from subsequent analyses if any of their intensity values fall in that range.

Our first approach was to truncate the normal distribution, similarly to our previous work [16]. In our previous work we employed this as a pruning operation to speed up the runtime of the forward algorithm, but reusing this technique to ignore highly unlikely data showed it was ineffective in this context. Instead, we considered two additional approaches, which we use together for greater data integrity.

First, we preprocess our data, either fitting or using expert judgment to determine ranges of fluorescence intensity values that are reasonable for the expected fluorophore counts. An example of this process is shown in **Figure 6**. This allows us to filter out reads containing improbable intensities that are most likely indicative of a contaminant unrelated to fluorophore signals.

Second, we determine, again through either fitting or expert judgment, the mu and sigma values of every fluorophore color channel in our reads. Doing this through Baum-Welch is unnecessary, as while Baum-Welch is helpful to us in determining parameter values that may interact in unexpected ways, fluorophore intensity distributions should be completely independent of the other parameters so long as we can effectively deconvolute varying fluorophore counts in our analysis. An example of mu, sigma, and background-mu estimation is shown in **Figure 6B**. We then hold these constant while fitting the other parameters.

Background sigma (the dispersion of the zero-count distribution) is computed automatically through a different mechanism. During signal processing with *sigproc_v2*, the image processing routine in the fluorosequencing data analysis package *plaster* (e.g. see [16]), signal is extracted from the central pixels of a peak, and a local background value is subtracted from the signal. The local background is computed by averaging the values of pixels just outside those used to compute the signal. The median value of these local background values is then used as the background sigma. We have empirically found this to work well in practice.

### A straightforward alternative method of parameter estimation using Powell’s method

As a second method of parameter estimation and an independent check on the HMM-based approach, we considered a relatively simple alternative, in which we minimize an objective function: every read in the data is first reduced to a dye track, which is an integer approximation of the original read, where the integers represent the best estimates of the numbers of fluorophores of each color at each time step. Given a parameter estimation, we can then simulate dye tracks, and compute the root-mean square error (RMSE) between the counts of each dye track in the original data and the simulated result.

To minimize the RMSE, we used Powell’s method [18], which allows us to optimize a function for which the derivatives cannot be computed. It proceeds by, along one dimension (parameter) at a time, finding the minimum, and repeating this for each dimension (parameter) until desired convergence criteria are achieved.

Empirically, we found that Powell’s method often halted in local minima. Thus, we first used the DIRECT global optimization method (**Figure 7A**) [19,20] to identify suitable initial conditions for Powell’s method (**Figure 7B**). DIRECT proceeds by repeatedly trying two test points above and below a given starting point along a particular dimension, and choosing the best among those and the original point. It then chooses another dimension and repeats until desired specificity of the result is achieved.

**Figure 7.**
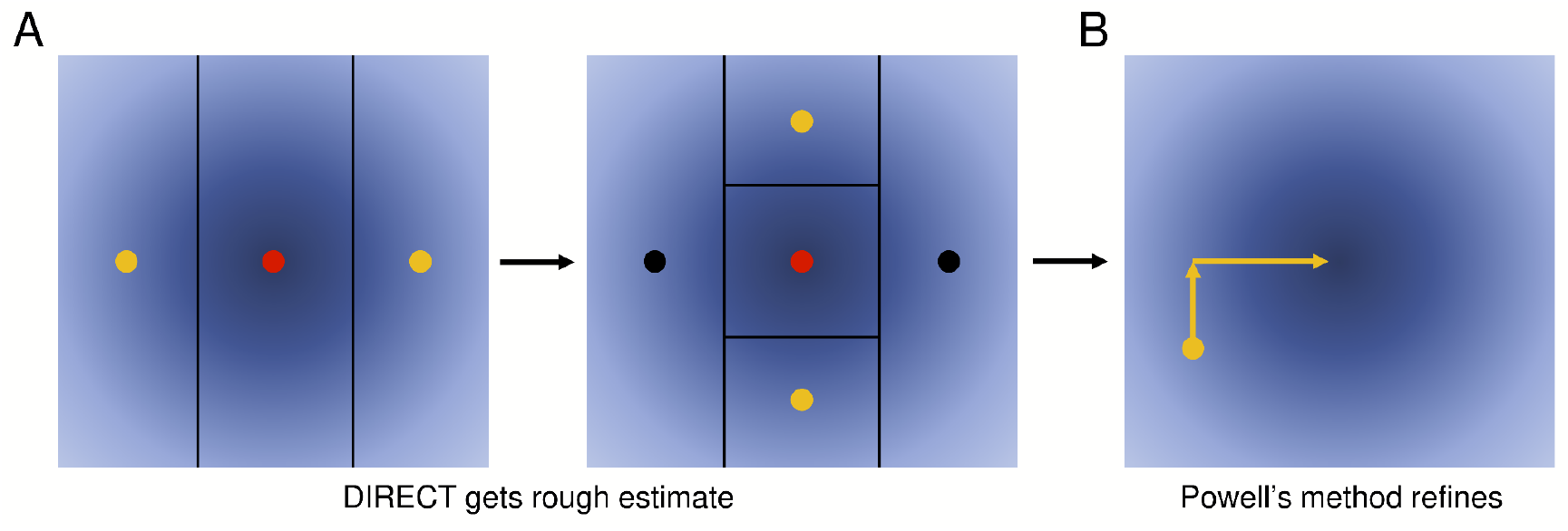
Illustration of DIRECT and Powell’s method. As an alternative to Baum-Welch, we also explored a more general purpose approach. (A) We first apply DIRECT to rapidly identify a region of the parameter space that is likely to contain the global optimum (red dots). DIRECT proceeds by iteratively comparing three points and using these results to further subdivide the search space, as shown. (B) We then apply Powell’s method, which iteratively minimizes the objective function by changing one variable at a time.

We found this combination of DIRECT and Powell’s method to be generally effective as an alternative to Baum-Welch, with the caveat of first having to independently estimate dye tracks, which may introduce changes relative to those considered during Baum-Welch.

### Overdetermined systems for low fluorophore counts

For peptides with only one label, many of the parameters are indistinguishable or uninterpretable. In particular, fluorophore destruction and peptide detachment are visibly equivalent when there is only one fluorophore; a fluorophore lost in the middle of a run could just as easily be a result of either effect. To account for this, when fitting parameters on peptides with one fluorophore, we fix the detachment rate to zero.

Another problematic parameter with peptides of only one label is the missing fluorophore rate, as it is not possible to detect peptides if all fluorophores are missing. When there are two labels on the peptide, we can correct for missing fluorophore biases as described above. But with one, as no peptides with missing fluorophores can be detected, we have no data to ascertain this probability. We therefore hold the missing fluorophore rate to zero in this case as well.

Initial and cyclic blocking rates are difficult to distinguish as well. If the Edman failure rate is allowed to move freely, then the initial and cyclic blocking rates are interchangeable, following an involved formulation:

Consider a peptide with only one fluorophore on the amino acid at position *r*. It can be described by an Edman failure rate of *e*, an initial block rate of *b*, and a cyclic block rate of *c*.

Consider an alternative parameterization of the data, where κ_*i*_ for *i* ≥ 0 represents the observed probability of the fluorophore’s last cycle being cycle *r* + *i*. We can relate these two interpretations with the formula:

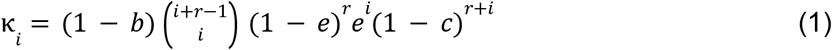

The probability at each cycle can be solved for in terms of the preceding cycle:

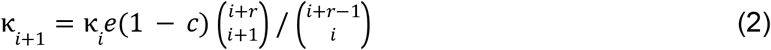

This simplifies to:

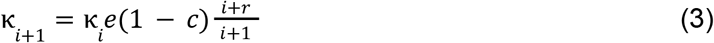

Then the quantity *e*(1 − *c*) effectively acts together as a single constant, which we can solve for using the ratio between κ_*i*+1_ and κ_*i*_ for some *i*. A choice of *e*(1 − *c*) uniquely determines all such ratios, therefore no number of equations for variables κ_*i*_ provides any additional information. We can however consider, instead of the ratio, one κ_*i*_ on its own. This would allow us to solve for *b* given a choice of *e* and *c*. We then have two equations and three unknowns; the system is underdetermined. Because of this, we hold the cyclic blocking rate to zero when only one fluorophore is present on the peptide.

### Bootstrapping

As knowing the precision of a result is important, we provided bootstrapping functionality for both methods by sampling the original data with replacement to create a dataset of the same size. This process can be repeated a user-defined number of times. We also provide, for both methods, the ability to construct a confidence interval using the percentile method of a user-defined size.

### Peptide notation in figures

In subsequent figures and legends, we indicate the amino acid sequences of the peptides being studied as single-letter amino acid codes. For simplicity in this paper, most of the peptides analyzed only have their lysines labeled, and lysines (denoted ‘K’) are followed by an asterisk, which indicates that they are labeled. Some peptides contained azido-lysine instead of lysine, denoted ‘{azK}’ which is, like lysine, followed by an asterisk indicating labeling. We notate the status of the N-terminus prior to sequencing, which is either “fmoc-” referring to the fluorenylmethoxycarbonyl protecting group (which is removed prior to sequencing), “ac-” referring to an acetylated N-terminal residue (incorporated to intentionally stop Edman degradation from occurring), or “NH_2_-” referring to the ordinary unaltered N-terminal state. For simplicity, we also truncate all sequences to the last lysine or azidolysine, as we do not expect amino acids following the last labeled residue to have an impact on the sequencing parameter estimation. We end the sequence notation with a “|” to indicate this omission.

### Experimental fluorosequencing control datasets

We fluorosequenced a number of purified peptides in order to assess parameter estimation on real experimental datasets, including control experiments in which we introduced modifications to the peptides or experimental workflow intended to impact specific error rates. All peptides were synthesized by Genscript and labeled without further purification. The labeled peptides were purified by HPLC and their mass and purity confirmed by LC/MS or SDS page gel purification as described in full in [21]. Briefly, the dye Atto643 was covalently attached to lysine or azido-lysine as appropriate by labeling directly with Atto643 NHS ester or with Atto643 conjugated *via* a polyethylene glycol (PEG)/polyproline tether (termed a Promer) to mitigate dye-dye interactions as in [21]. Peptides were fluorosequenced with a minimum of 40,000 reads using the same sequencing workflow and Total Internal Reflection Fluorescence (TIRF) microscopes denoted Systems A & B in [21].

## Results

### Estimating parameters from simulated datasets as a first test of the algorithms

We evaluated the performance of our modified Baum-Welch algorithm and our DIRECT/Powell’s parameter estimation method by analyzing a number of fluorosequencing datasets, some simulated and subsequently others from actual experimental datasets, including ones with controlled manipulation of experimental parameters to test the sensitivity and concordance of the parameter estimators to such effects. We first considered fully simulated datasets, in which the true values of the parameters were known precisely and the (simulated) sequencing process adhered perfectly to the sequencing process implicit in the HMM or Powell’s analysis.

As a first test, we used a previously developed Monte Carlo simulation of fluorosequencing [16] to generate simulated fluorosequencing reads in order to validate the parameter estimators against synthetic datasets with precisely known parameters. With these datasets and known parameters in hand, we then tested the parameter estimators for their accuracies. **Figure 8** plots results for both fitting tools for a two-fluorophore peptide, allowing fitting of all parameters.

**Figure 8.**
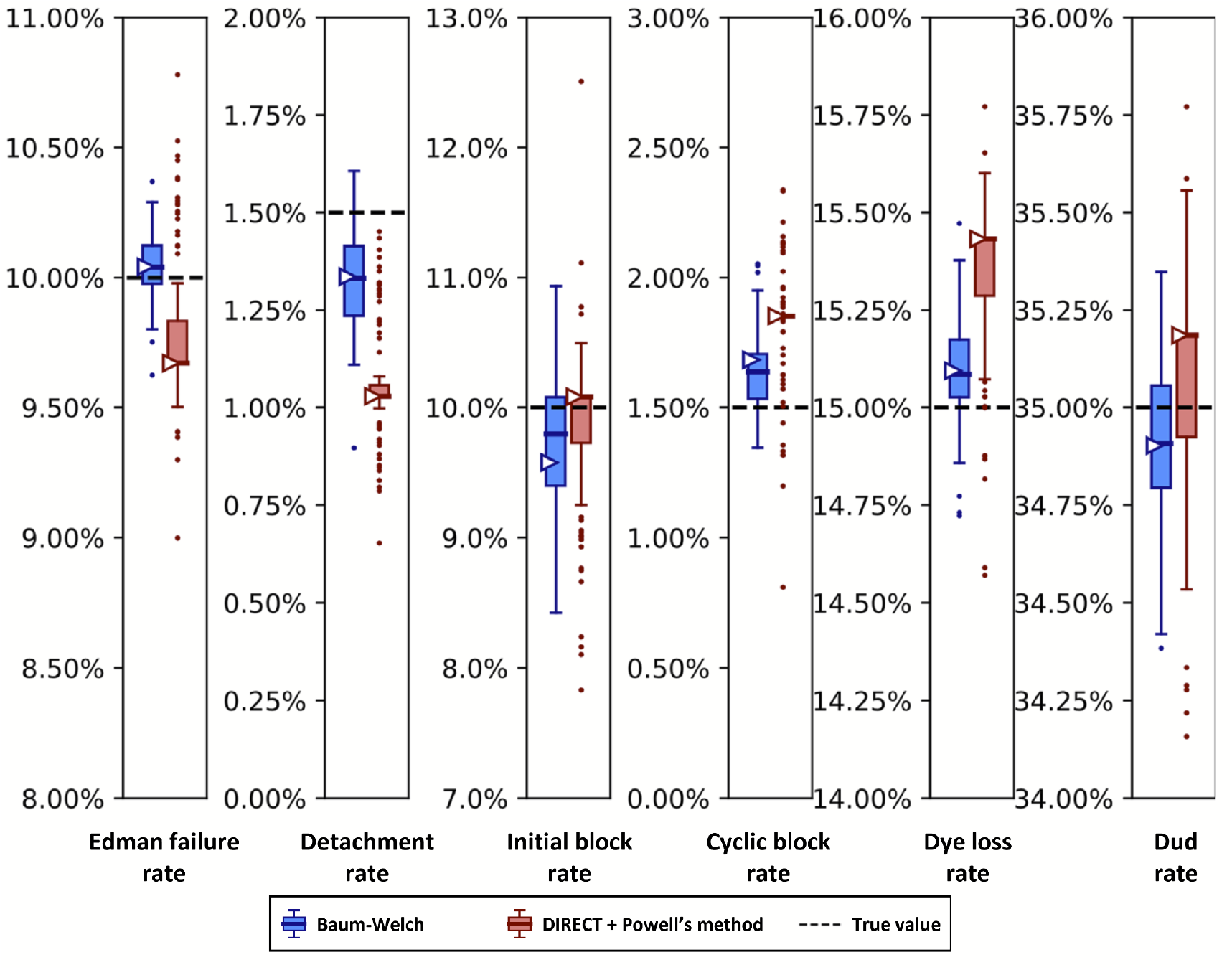
The Baum-Welch and DIRECT + Powell’s methods agree within 0.5% error on simulated data. Synthetic fluorosequencing reads were generated for peptide NH2-G{azK}*AG{azK}*|, and the simulated dataset was bootstrapped by subsampling with replacement the same number of reads 100 times. Parameters were estimated for each of these bootstrapped subsamples with both methods as described in the text. The box-and-whisker plots represent the distributions of these results, plotting 1st and 3rd quartile +/-max/min observation within 1.5 interquartile range (IQR). Results outside 1.5 times the IQR are considered outliers and are plotted as points. The right facing triangles mark the parameter estimate found if using the non-bootstrapped original data. The dashed black lines indicate the target value that was used in simulation.

Using both the Baum-Welch and DIRECT followed by Powell’s method allowed us to explore potential strengths and weaknesses in both approaches. We saw that our Baum-Welch based approach was able to consistently include the true value within 1.5 times the interquartile range (IQR) of the bootstrapped parameter estimates. For the same synthetic dataset, using DIRECT followed by Powell’s method appeared somewhat less reliable -the detachment rate was particularly poorly fit, with the target value lying outside of the range of even the outliers, while the cyclic block rate and the dye loss rate were outside of 1.5 times the IQR but within the more extreme maxima and minima of the outliers. Nevertheless, both methods were moderately close to the target value in most cases, and in all cases, both their primary and median bootstrapped estimates agreed with the true value within 0.5% absolute error.

In order to better understand the effects of variably sized datasets on the size of the confidence intervals given by the Baum-Welch based approach, we next explored this relationship specifically. We anticipated that the scale of the distribution of results would scale with the inverse of the square root of the number of reads in the dataset, as this is the behavior of most distributions in the limit as the number of datapoints rises. However given the complexity of the process being modeled, we thought it wise to verify this by simulation. In **Figure 9**, we indeed find that the dispersion of the parameter distributions shrinks in the predicted manner, generally converging nicely when analyzing more than 10,000 to 100,000 reads.

**Figure 9.**
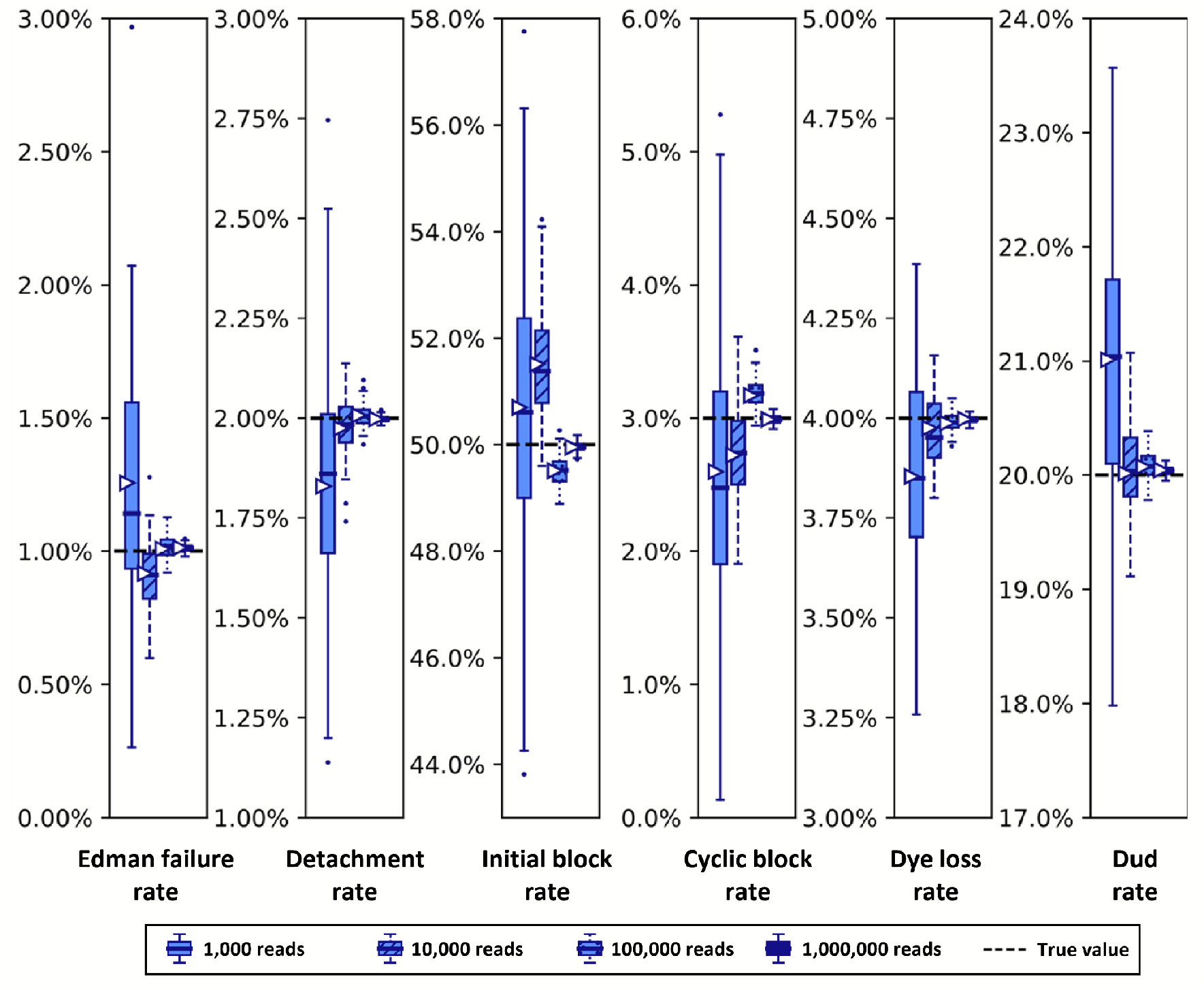
Simulated datasets with more reads exhibit tighter distributions of parameter estimates. Synthetic fluorosequencing reads were created for peptide NH2-G{azK}*AG{azK}*|, simulated the numbers of reads indicated in the graphical legend. These datasets were bootstrapped by subsampling with replacement the same number of datapoints as in the associated simulated dataset, with 100 bootstrapping rounds, and estimating parameters for each of these bootstrapped subsamples with both methods as described in the text. Box-and-whisker plots are defined as in **Figure 8**.

### Parameter estimation from experimental fluorosequencing datasets

While reproducing the parameters used to generate synthetic data demonstrates the basic validity of the algorithms, they must be able to draw robust estimates from real experimental datasets. We thus collected a series of experimental sequencing datasets designed to intentionally manipulate isolated (where possible) error rates and tested the parameter estimators on these datasets. In all cases, control peptides of known sequence were synthesized, purified, and analyzed, labeling the peptides’ lysines (or azido-lysines, as appropriate) with Atto643 dyes on Promer linkers as in [21]. Labeled peptides were then fluorosequenced, collecting a minimum of 30,000 reads. The full details and all experimental procedures and additional control experiments are reported in [21].

Importantly, sequencing a known peptide labeled with two fluorophores allows us to fit all six free parameters simultaneously. We demonstrate this for both fitting methods with two independently collected experimental datasets in **Figure 10**. In general, the methods estimated parameters within a few percent of each other. However, we noted a tendency for DIRECT followed by Powell’s method to give results on the original dataset (triangle markers) distant from the median results of the bootstrapped data (box plot midlines), in some cases even outside the first-to-third quartile range. Its distributions also tended to be much broader than those given by Baum-Welch, suggesting less confidence in its estimates. However an alternative interpretation could be considered: the wider distribution of DIRECT followed by Powell’s method allowed for overlapping distributions across the replicate datasets in every case except for the missing fluorophore (dud) rate, while Baum-Welch produced distributions that fail to overlap completely for not only the missing fluorophore rate, but also the initial block rate, and the Edman failure rate had only the slightest overlap. Thus, DIRECT+Powell’s appeared more consistent with the scale of parameter variation observed across experimental replicates.

**Figure 10.**
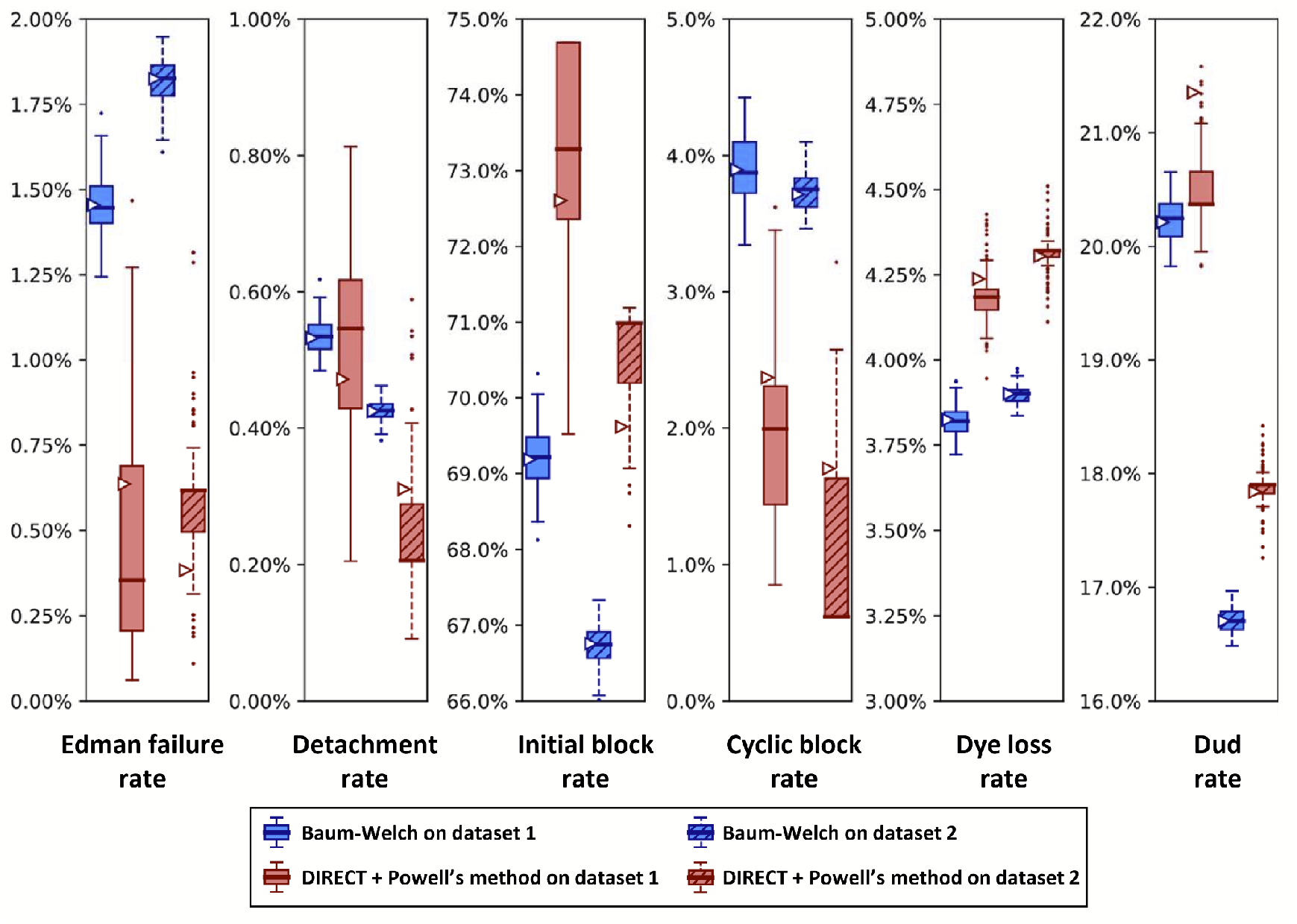
A comparison of Baum-Welch and DIRECT + Powell’s method on experimental sequencing data for a two-fluorophore peptide shows general agreement between the methods. Fluorosequencing datasets for peptide NH2-G{azK}*AG{azK}*| were collected one day apart by the same researcher on the 21st (dataset 1 with 40,181 reads) and the 22nd (dataset 2 with 71,823 reads) of November 2022. The original datasets were bootstrapped by subsampling with replacement the same number of datapoints as in the original dataset 100 times, and fitting on each of these bootstrapped subsamples with both methods as described in the text. Box-and-whisker plots are defined as in **Figure 8**.

We also observed an unusual tendency in **Figures 8 and 10** for DIRECT with Powell’s method to produce distributions of fits from bootstrapped data with large numbers of outliers, with medians sitting on top of the first or third quartile, and in one case (cyclic block rate in **Figure 10**) the median even appears to be identical to the minimum. We do not have an explanation for this phenomenon.

In the course of other studies, we noticed varying Edman cleavage efficiencies at proline residues, which had been purported to sequence less efficiently in historic Edman degradation experiments [22–25]. Initial fluorosequencing studies saw little effect stemming from proline [13], but subsequent studies showed less efficient cleavage, an effect we were able to trace to the length of incubation time in trifluoroacetic acid (TFA) [21], a key reagent for Edman degradation whose usage had been reduced while optimizing the sequencing workflow. This effect thus provides a means to experimentally manipulate the Edman failure rate and test the response of the parameter estimators.

Figure 11. presents such an analysis, plotting the results of analyzing two independently collected fluorosequencing datasets for the same proline-containing peptide with a single fluorophore in the third position. Results for Baum-Welch differed considerably, while the DIRECT with Powell’s method remained more consistent. In particular, the Baum-Welch method’s discrepancy in the Edman failure rate between the two experiments, predicting values of approximately either 12% or 7%, is concerning, as while it is possible that the true value did vary by this amount, the DIRECT/Powell’s method shows higher concordance between the experiments.

**Figure 11.**
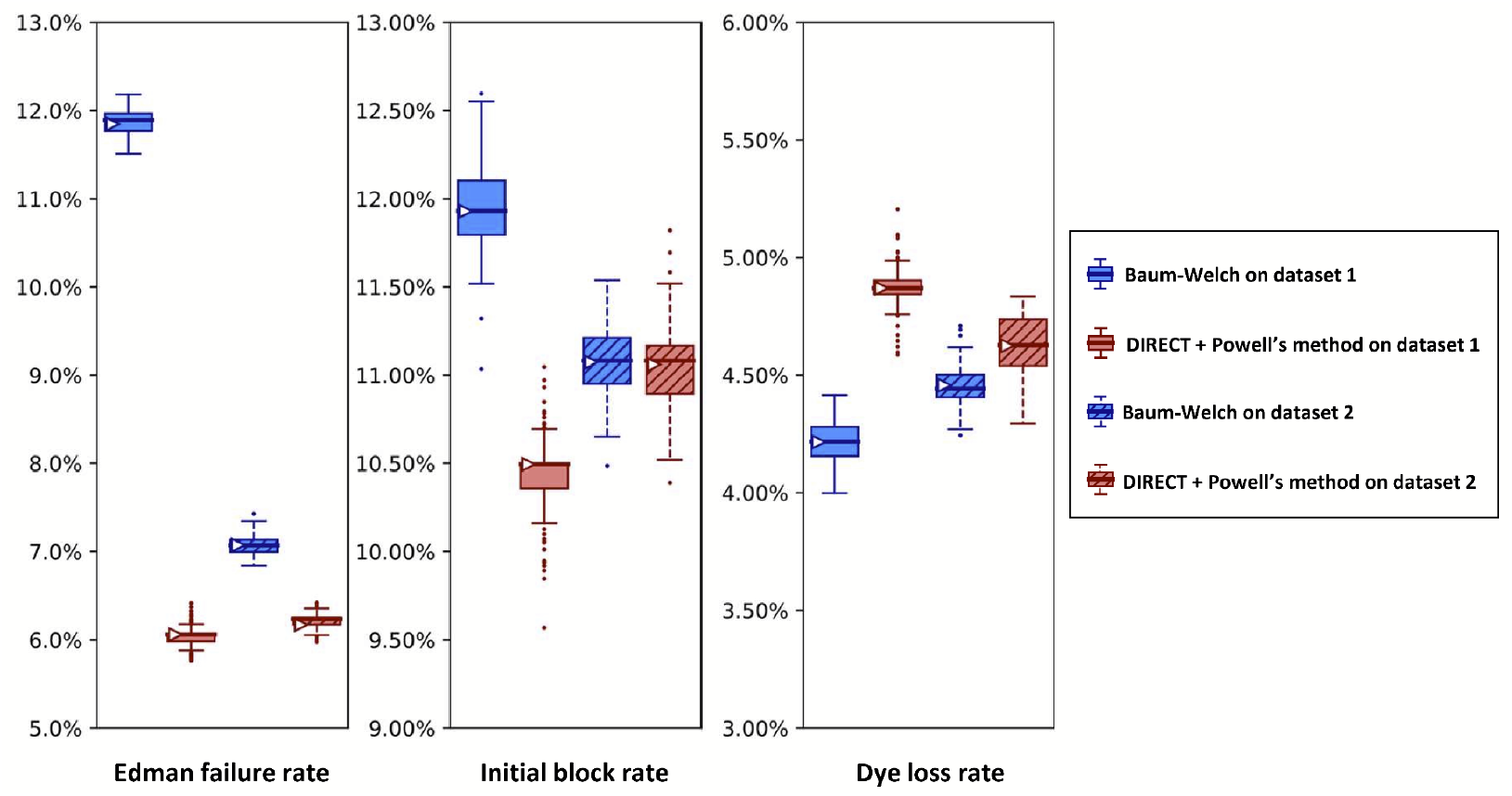
A higher Edman failure rate is observed for a proline-containing peptide, with general agreement between the estimation methods. Here, we analyze two experimental fluorosequencing datasets for peptide fmoc-APK*| collected by the same researcher on the 11th (dataset 1 with 27,783 reads) and 28th (dataset 2 with 34,380 reads) of November 2022. The original datasets were bootstrapped by subsampling with replacement the same number of datapoints as in the original dataset 100 times, and fitting on each of these bootstrapped subsamples with both methods as described in the text. Box-and-whisker plots are defined as in **Figure 8**.

As a comparison with the proline containing peptide in **Figure 11**, in **Figure 12** we present results for a peptide that has no prolines. As expected, the no-proline case has a much higher Edman cleavage efficiency (1 -Edman failure rate) than for the case of the proline-containing peptide. We also see a significantly higher initial block rate and dye loss rate, the former of which may relate to less efficient cleavage at the glycine residue.

**Figure 12.**
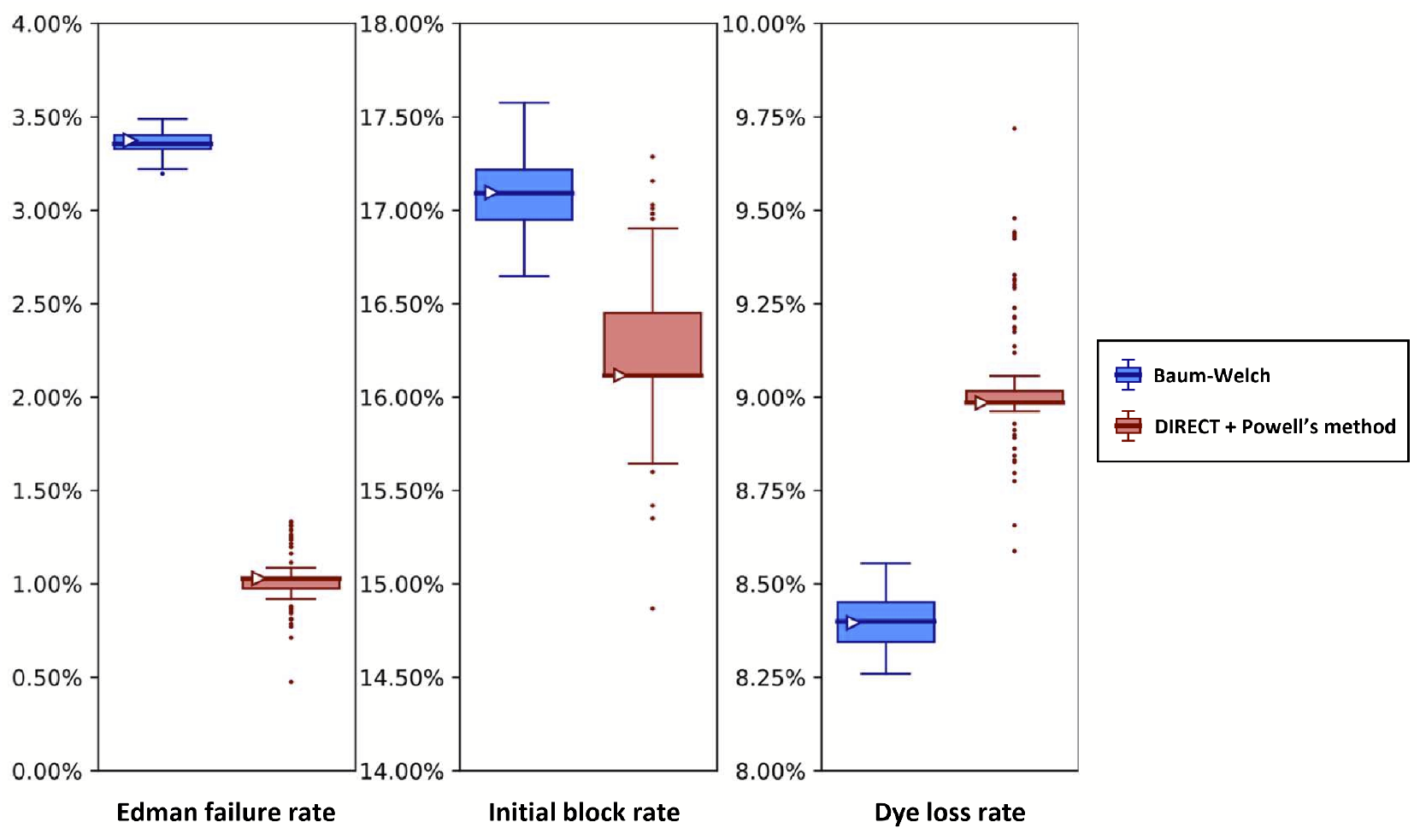
Analysis of experimental fluorosequencing data from a peptide similar to that in Figure 11 but containing no proline residues exhibits lower Edman failure rates and shows agreement between the methods. The peptide fmoc-GAK*| was sequenced on December 1st, 2022, and the original dataset of 67,936 reads was bootstrapped by subsampling with replacement the same number of datapoints as in the original dataset 100 times, and fitting on each of these bootstrapped subsamples with both methods as described in the text. Box-and-whisker plots are defined as in **Figure 8**.

We would expect an acetylated, or blocked, peptide to be fit with either a very high Edman failure rate or initial block rate, or perhaps both. **Figure 13** confirmed this prediction, with initial block rates strictly above 91% for all bootstrapped results with either dataset and either fitting technique. While these results raise our confidence in both techniques, they are lower than anticipated. We suspect that contaminants captured by accident in this dataset are responsible for this behavior. We also conjecture that the 40% range of discrepancies between datasets in the Edman failure rates arose because of low signal and increased sensitivity to contaminants due to the high initial block rate.

**Figure 13.**
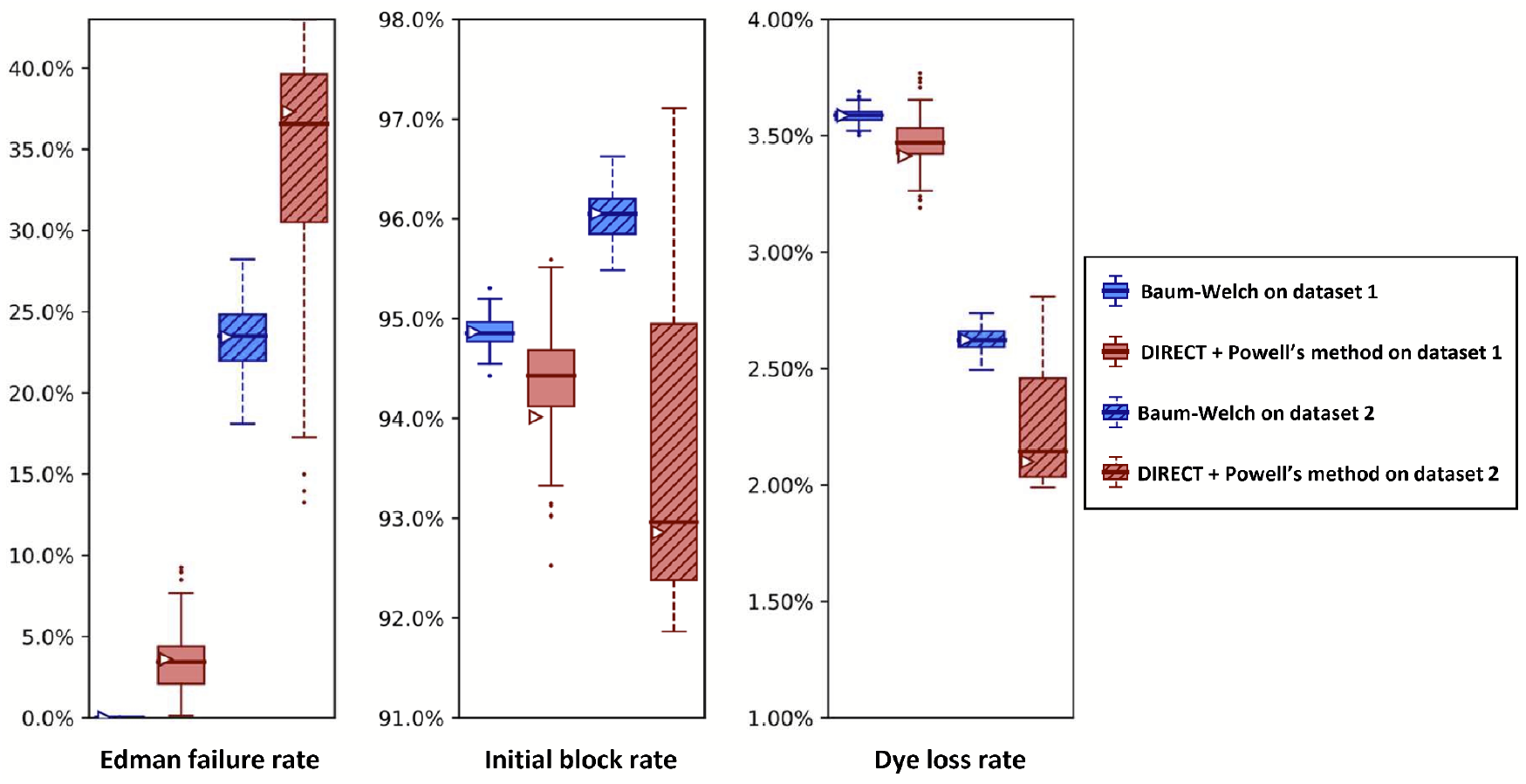
Both parameter estimators correctly recognize high initial block rates (>91%) in an intentionally N-terminally blocked peptide. Here, we examine two independent fluorosequencing datasets for ac-A{azK}*|, an N-terminally acetylated peptide, *i*.*e*. one for which the N-terminus is covalently blocked and not sequenceable by Edman chemistry. The datasets were collected on the 15th (dataset 1 with 43,606 reads) and 17th (dataset 2 with 42,417 reads) of November 2022 by the same researcher. The original datasets were bootstrapped by subsampling with replacement the same number of reads as in the original dataset 100 times, and fitting on each of these bootstrapped subsamples with both methods as described in the text. Box-and-whisker plots are defined as in **Figure 8**.

In order to test whether other parameters were being correctly estimated, we also manipulated the incubation time in trifluoroacetic acid (TFA), which should directly relate to Edman failure rates, for each of 6 independent peptides. TFA is a key reagent for Edman degradation (**Figure 1**), but also one known to inactivate dyes with sufficiently long incubations [13]. In our initial fluorosequencing paper, we performed 15 or 30 minute incubations [13], but subsequent tests revealed that incubation times could be shortened with little loss of cleavage efficiency [21]. Thus, we wished to test the minimum TFA incubation times and explore the intrinsic tradeoff between cleavage efficiency and dye loss rates.

Figure 14. plots how the various estimated error rates changed on experimental fluorosequencing data collected with 4 different TFA incubation times during the Edman degradation steps. As expected, reducing TFA incubation time to near zero increases the Edman failure rate for all peptides studied, but with as little as 4-6 minutes, all peptides with the exception of the proline containing peptide exhibit low Edman failure rates, hence reasonably efficient cleavage efficiency (**Figure 14A**). The effect of longer TFA incubation time on the initial N-terminal blocking rate is less clear, with large effects primarily noted on the proline containing peptide, suggesting there may be some tradeoff by the parameter estimators in assigning weight to the Edman cycling versus initial block in that specific case (**Figure 14B**). Also as expected, longer exposure to TFA results in higher dye loss rates (**Figure 14C**), consistent with our prior observations.

**Figure 14.**
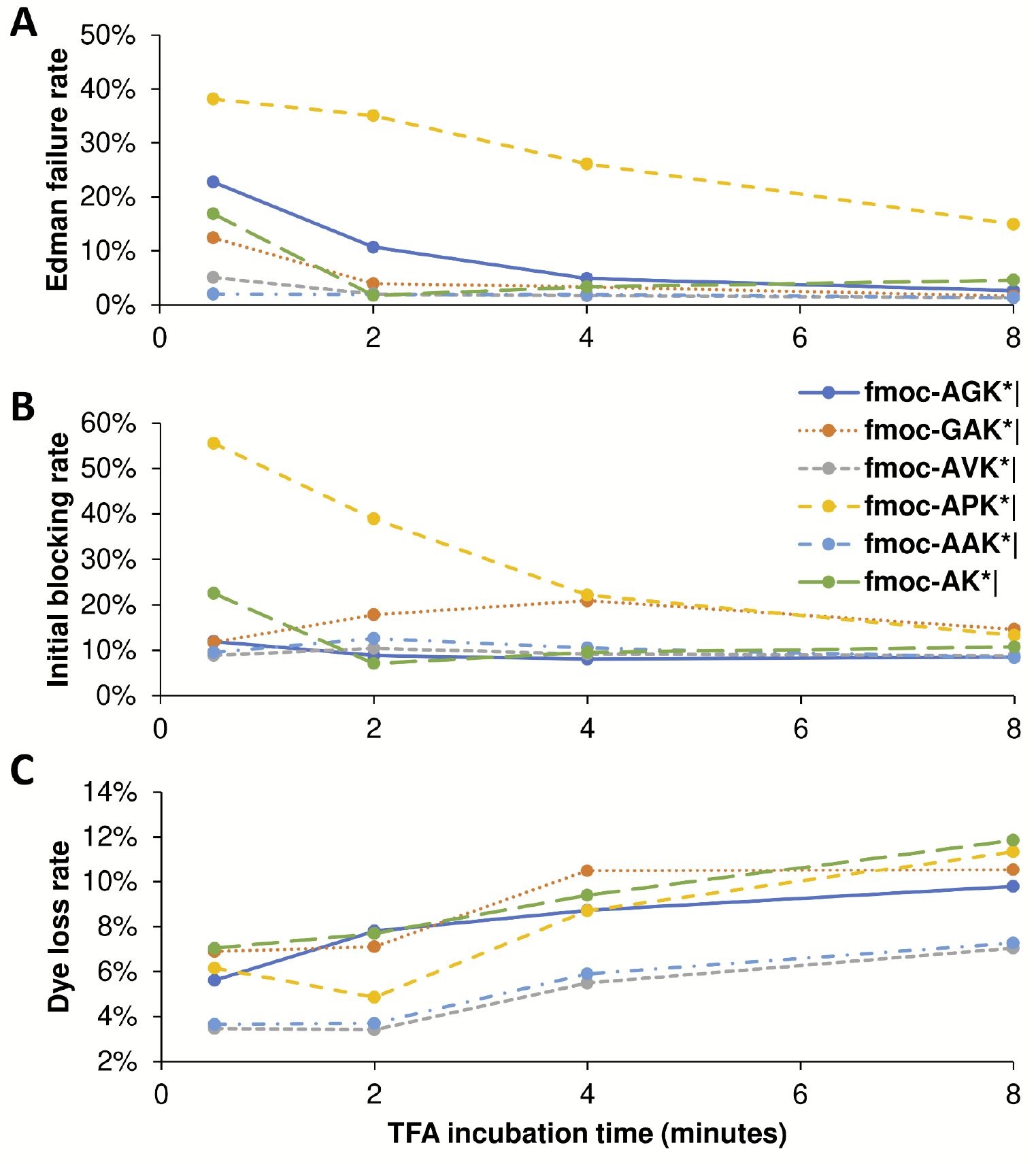
Longer TFA incubation times reduce the Edman failure rate but increase the dye loss rate. We tested minimum times required for TFA incubation by sequencing six different peptides with four different lengths of time of TFA exposure. Every dataset used in these plots has at least 34,000 reads, and all but three have more than 50,000 (see Zenodo repository for precise counts). (A) The Edman failure rate decreases with longer TFA incubation time, which matches theoretical expectations. (B) The initial block rate does not follow a clear pattern. (C) The dye loss rate goes up with longer TFA incubation time, which matches theoretical expectations.

## Discussion

A reason to use two parameter estimation methods was to measure their agreement on real data; in **Figure 10** we unfortunately see wide discrepancies between the two methods. This was not observed for the simulated data in **Figure 8**. We observed less severe modeling discrepancies for the real data plotted in **Figures 11 -13** as well. We believe these discrepancies are a result of the contaminants and other unknown factors that are present in data. This could cause slightly incorrect results for both models, but they may react differently in the case of an incomplete model.

Some of the more unusual results may be caused by errors in sample preparation or data collection. Across all real datasets (**Figures 10 through 14**), we observe variations in the initial block rate that do not follow a clear or consistent pattern. We suspect that some step in sample preparation or data collection is ultimately responsible, but we have as yet been unable to determine the root cause.

For other parameters, such as the dye loss rate, there are small but statistically significant fluctuations between experiments, not caused by modeling differences between the methods. As with the larger discrepancies in the initial block rate, inconsistent sample preparation or data-collection practices could be responsible. However other interpretations are possible. Indeed, some amount of inconsistency should be expected, as any number of factors can change, including variations in reagents or imaging quality. It is also possible that our data analysis practices are at fault. We have observed random fluctuations such that all intensity values may be a small amount brighter or dimmer at each cycle of our experiment. Applying the same distributional cut-offs and distributions of intensity across all time steps may then introduce biases in the data that is kept and in how it is analyzed, resulting in small fluctuations for multiple parameters. This is a possible avenue of future exploration.

We can also ask which of our methods is better. With a clear winner between Baum-Welch and Powell’s method, we could reduce future explorations to the better method. Baum-Welch provides more accurate results for simulated data; in **Figure 8** both the estimates on the original data and the medians of the bootstrapping results are better for five out of the six parameters with the Baum-Welch algorithm than with the Powell’s method based results.

Baum-Welch also tends to have tighter confidence intervals and more consistent distributional shapes than what we see with Powell’s method. Baum-Welch mostly appears to be the better technique, but questions remain, in particular the discrepancy between the two replicates shown in **Figure 11** for the Edman failure rate for Baum-Welch but not for Powell’s method.

Finally, our new parameter estimation techniques prompted us to compare our experimental results to previous, though less precise, results of fluorosequencing in earlier publications [13]. A number of advancements have been made which have improved these error rates [21]. In 2018 the Edman failure rate was measured at around 6%. Edman failure rates now appear to be around 1% or 2% for most residues and for much shorter TFA incubation times, following extensive optimization of the reaction conditions [21]. A consequence of the shorter TFA incubations, however, is that error rates at prolines remain higher, as in **Figure 11**. Dye loss rates and peptide detachment rates were estimated at 5% in 2018; while current results suggest an unchanged peptide detachment rate, the dye loss rate has been brought below 1%, perhaps to 0.5%.The missing fluorophore rate appears to have risen from 7% to 20% in this time-frame, and experiments may be needed to investigate the source of this rate.

## Conclusions

We have developed and tested two computational approaches for estimating key parameters governing protein fluorosequencing experiments, which should provide guidance for refining experimental protocols as well as improve the quality of peptide classification performed on the resulting datasets. One approach introduces modifications to the Baum-Welch algorithm that allow it to better isolate sources of error from each other and to then compute weighted MLEs at each step of the Baum-Welch Expectation Maximization process. The other approach minimizes the RMSE of the dye-track counts between simulations from the parameters and the provided dataset. Both methods seem promising and are generally concordant, and thus provide important checks on each other. We anticipate these approaches will be generally useful for ongoing efforts at improving the fluorosequencing experimental pipeline, including better identifying and mitigating mis-sequenced peptides and fluorescent contaminants, as well as for developing improved computational methods for interpreting fluorosequencing datasets.

## Acknowledgements

All code and data are available: whatprot (Baum-Welch) is downloadable from https://github.com/marcottelab/whatprot;sigproc_v2 (image processing) and pfit_v1 (DIRECT + Powell’s) from https://github.com/marcottelab/robust-fluorosequencing-plaster; and fluorosequencing datasets are deposited on Zenodo at https://zenodo.org/record/8137155. The authors gratefully acknowledge Jagannath Swaminathan, Eric Anslyn, and Daniel Weaver for helpful guidance and discussion throughout the course of this project. M.B.S. acknowledges support from a Computational Sciences, Engineering, and Mathematics graduate program fellowship. E.M.M. acknowledges support from Erisyon, Inc., the National Institute of General Medical Sciences (R35GM122480), the National Institute of Child Health and Human Development (HD085901), and the Welch Foundation (F-1515).

## Appendix A Baum-Welch and modifications

### A1 -Baum-Welch (in generic form)

The Baum-Welch algorithm is a form of Expectation Maximization, which will iterate over solutions to reach a maximum likelihood estimate. While only convergence to a local maximum likelihood can be guaranteed, it has been found that for most applications Expectation Maximization is nevertheless highly effective. Like all Expectation Maximization techniques, in Baum-Welch, we assume a parameterization to start, which we use to compute probabilities of intermediate hidden values, which we then use to produce a new and better estimate of the parameters. For Baum-Welch, this means iteratively improving the transition and emission probabilities.

The Baum-Welch algorithm is built partly upon the forward-backward algorithm. For any HMM we have a transition matrix and an emission matrix which define the model. Let *X*_1:*T*_ represent the random variables which take one of *N* possible values over *T* time steps. Let *Y*_1:*T*_ represent Nthe random variables representing the emission distribution. Let *s* denote the step in our Baum-Welch iteration process, and θ^(*s*)^ represent our choice of parameters for step *s*, which includes our transition probabilities, emission distributions, and our initial conditions. Our transition matrix is given as:

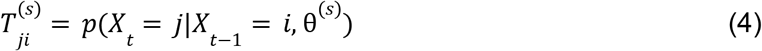

In most introductory descriptions of Baum-Welch, the emission values are from a discrete set, and an emission matrix is given as the probability of seeing a particular indexed output given a particular state. In this application, our emissions are floating point values, and we find it more clarifying to instead have a different emission matrix for every time step, and to think of it as a diagonal matrix where the values represent the probability density of the known observation at that time step. Letting *t* denote the time step, and *y* denote the true observed value at time *t*.

We define our emission matrix *O*^(*s,t*)^ as follows:

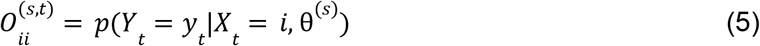

We need to run the forward algorithm. We define our cumulative forward probabilities as:

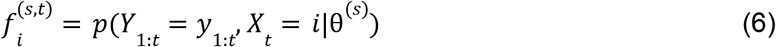

We can use dynamic programming to determine 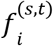 for every value of *i* and *t*, if given initial conditions 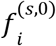 for all *i*. The equation is:

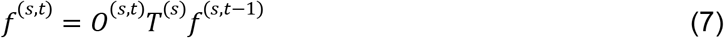

Or equivalently:

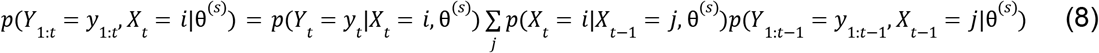

We also need to run the backward algorithm. We define the probabilities as:

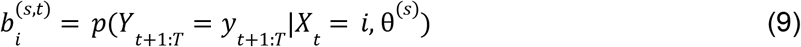

We can use dynamic programming to determine *b*_*i*_^(*t*)^ for every value of *i* and *t*. We initialize it with:

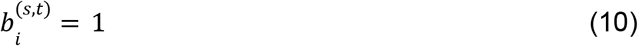

And then compute:

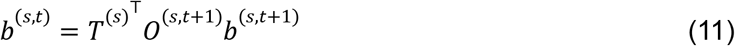

Or equivalently:

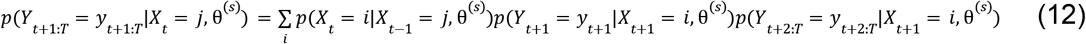

This gives us the capability to compute two sets of key intermediate values.

The first is the same as what is computed in the forward-backward algorithm, and represents the probability of being in a state at a given time given the entire sequence of observations. It’s defined as:

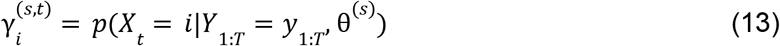

And computed with the formula:

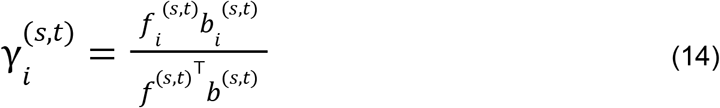

Or equivalently:

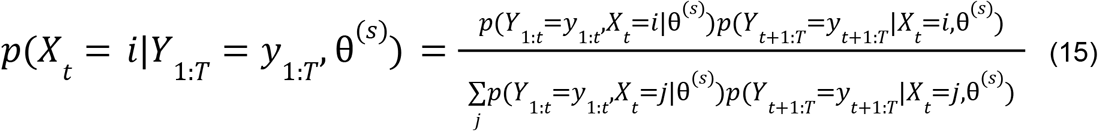

The next set of values represents the probability of a particular transition taking place between time step *t* and time step *t* + 1. We define it as:

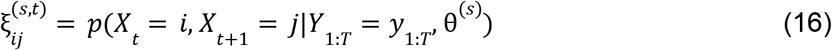

We can compute this with:

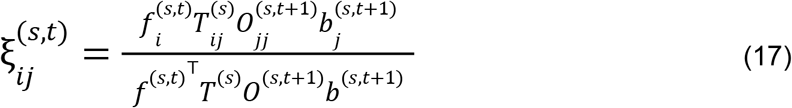

Or equivalently:

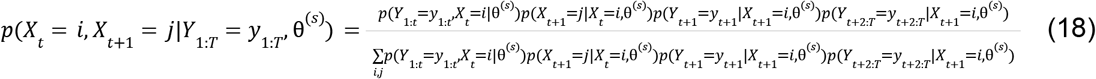

We then update our transition probabilities according to:

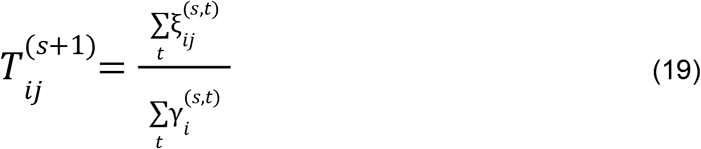

Or the equivalent expression:

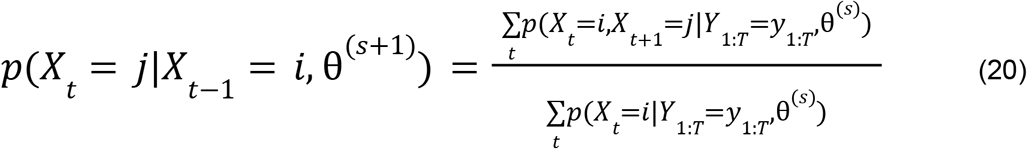

We also need to determine our new initial conditions. We solve for these as:

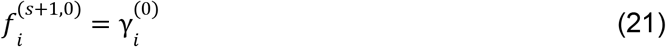

To update our emission probabilities, we do a weighted maximum likelihood estimate, using ξ to determine the appropriate weights. The form of the maximum likelihood estimate depends on the assumed shape of the output distribution, but if we assume each output distribution is normally distributed, where 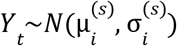 when *X*_*t*_ = *i*, we can compute μ^(*s*+1)^ and σ^(*s*+1)^as:

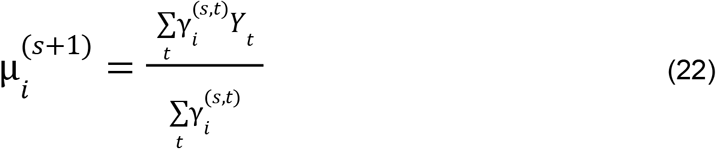

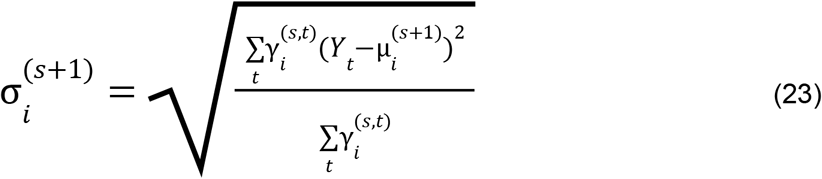

Or the equivalent expressions:

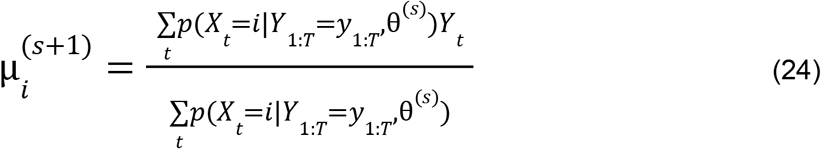

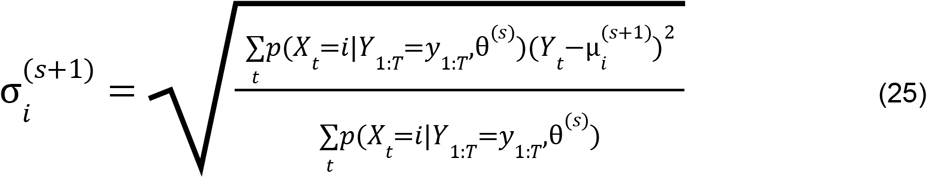

We can then compute a new emission matrix using these new parameter estimates:

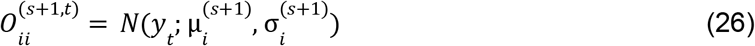

This can be trivially generalized to multiple sequences. Let *r* denote the index of a sequence, ranging from 1 to R. Then:

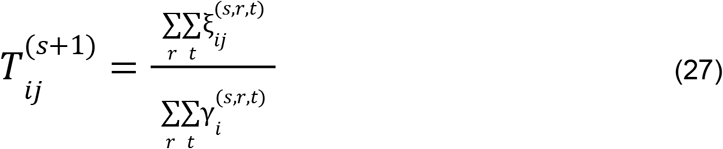

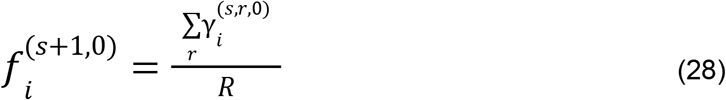

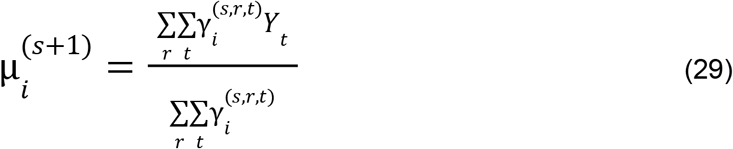

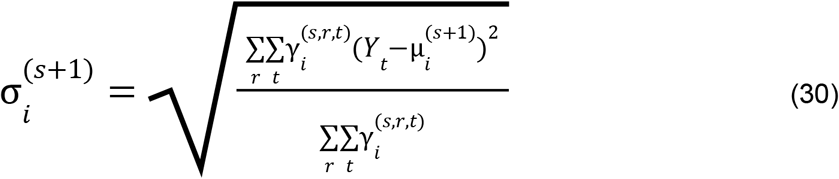

### A2 -Baum-Welch with isolation of error types

We now revisit Baum-Welch under the consideration of a factorization of the transition matrix, given as the product of *U* matrix factors:

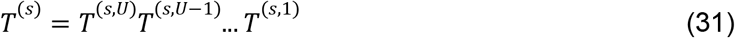

Then we find that:

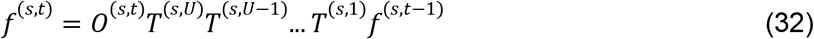

We can think of these factors as transition probabilities between intermediate states. We define random variables of the form *X*_*t,u*_ and letting *X*_*t*,0_ = *X*_*t*_. We find that:

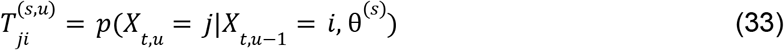

It is useful to instead track intermediate cumulative values in this formulation. Let:

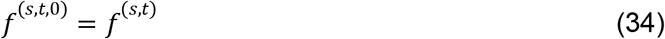

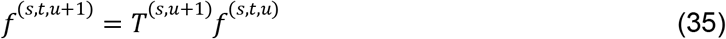

Then it follows that:

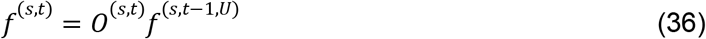

These values represent the probabilities of various states after sub-transitions, given as:

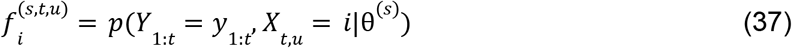

Through a similar re-engineering of the backwards recursion we get:

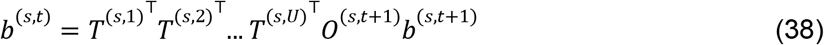

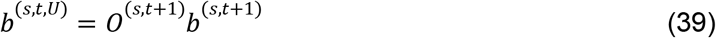

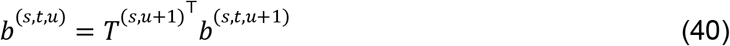

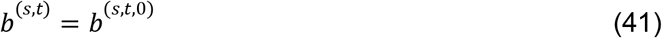

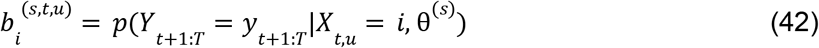

We then need to track the probabilities of these intermediate states and transitions. We find them to be:

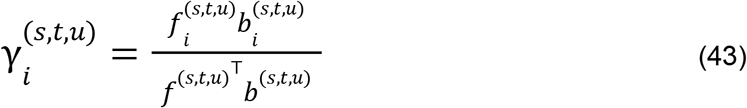

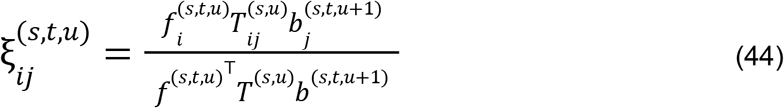

Noting that ξ^(*s,t,u*)^ is only defined when *u* < *U*, we then update our factored transition probabilities using the equation:

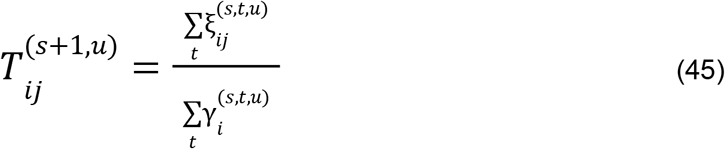

The initial conditions and emission matrices can be computed as before, if we note that:

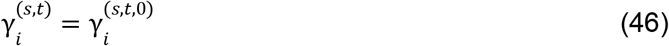

However, our initial conditions and emission probabilities have structure for fluorosequencing which will be helpful to define. In particular, our initial conditions are affected only by the missing fluorophore rate and the initial-blocking rate. We find it easiest to pretend we start with a perfectly labeled and non-blocked peptide, and then we apply a pre-transition which is different from the one used between emissions. Letting τ represent the pre-transition, we then use this pre-transition in the following manner:

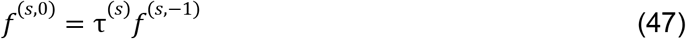

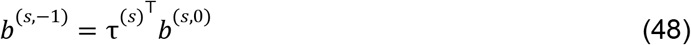

Letting 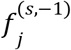 be 1 for the perfectly labeled and non-blocked state, and 0 everywhere else, while allowing the factorization τ^(*s*)^ = τ^(*s,V*)^τ^(*s,V*−1)^. τ^(*s*,1)^ as was done with *T*^(*s*)^ previously. The transition probabilities of τ^(*s*)^ can then be computed in manner just like that for *T*^(*s*)^:

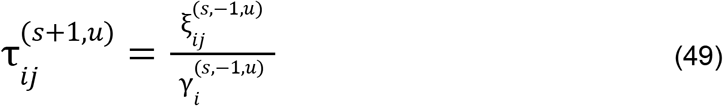

We also consider the handling of *O*^(*s,t*)^. In particular our output space is multidimensional, such that *Y*_*t*_ ∈ ℝ^*W*^, so we let *Y*_*t*_ = (*Y* _*t*,1_, *Y* _*t*,2_, ., *Y* _*t,W*_). If these components of the output are independent Nrandom variables, we can let:

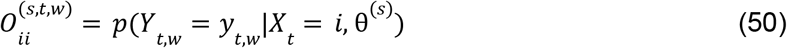

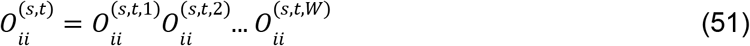

And we update our emission matrices as before. Assuming a normal distribution gives us:

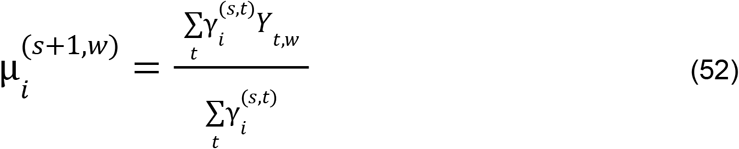

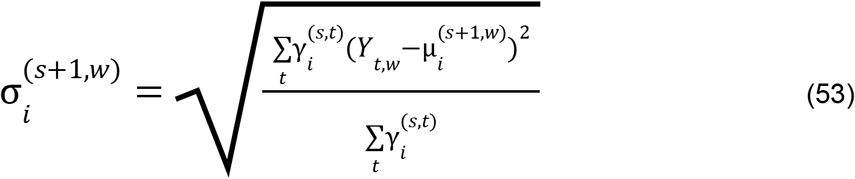

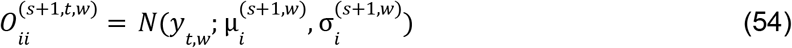

We can still generalize this to more sequences, using the generalized equations:

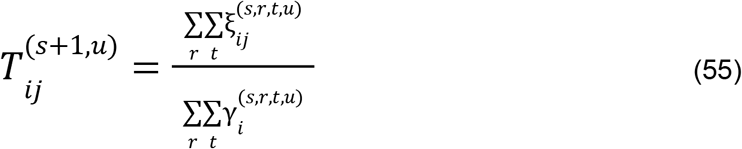

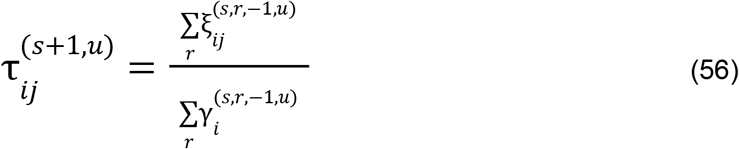

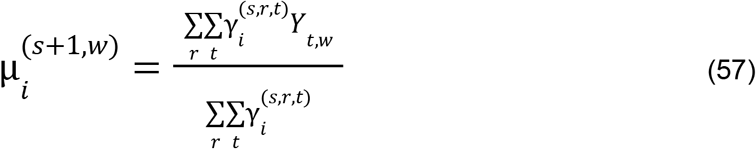

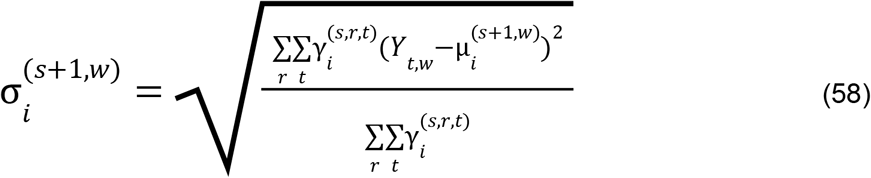

### A3 -Baum Welch with weighted Maximum Likelihood Estimation

Instead of getting transition probabilities, we may prefer to get a direct maximum likelihood estimate of a more fundamental parameter value that provides a more interpretable result. Furthermore by involving a stricter structure in our formulation, parameterized by less parameters, our results should be less prone to error from overfitting the data. We will achieve this by means of a technique resembling what is done to estimate the parameterization of the continuous distributions for the emission matrix.

For every transition factor indexed by *u*, we introduce a parameter *p*_*s*+1,*u*_, a function, *g* _*u*_ : ℕ → ℕ, a function *h*_*u*_ : ℕ × ℕ → ℕ, and a function ψ _*u*_ : ℝ → ℝ ^*N*×*N*^. The form of these functions depends on the form of the random variable being modeled by the transition factor, and we will describe this in more detail below. For now we define:

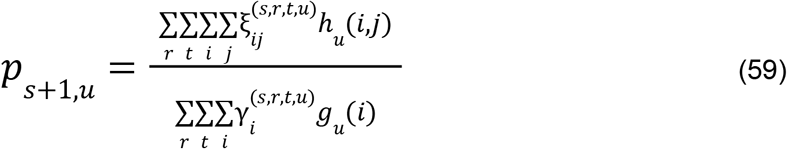

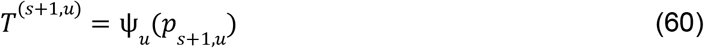

Similarly for our initial conditions we let:

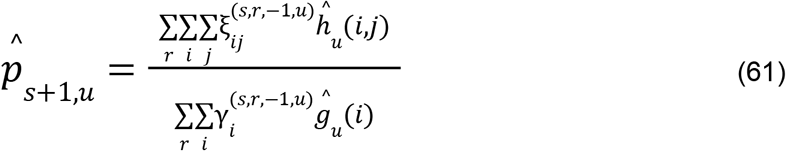

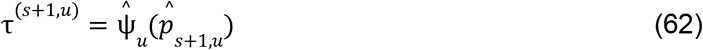

Now we can define our factor specific functions.

#### Detachment rate

We start with the detachment rate because it is the easiest to describe. It is defined by a Bernoulli random variable, which determines the rate at which a peptide enters a detached state instead of remaining unchanged.

We can only measure the detachment rate from states with remaining amino acids. We therefore set *g* _*u*_ (*i*) to 1 for those states, and 0 for the remaining state. To pick up the transition rate into these states, we set *h* _*u*_ (*i, j*) to 1 if *i* has remaining amino acids *and j* indicates the detached state. We set it to 0 for all other combinations of *i* and *j*, which should never occur as they should have been forbidden in the definition of the transition factor in the previous iteration of Baum-Welch, *T*^(*s,u*)^.

This is a weighted maximum likelihood estimate of a Bernoulli random variable. We use the resulting value of *p*_*s*+1,*u*_ to define the corresponding transition factor. ψ _*u*_ then defines 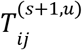 to be *p*_*s*+1,*u*_ when *i* has remaining amino acids *and j* indicates the detached state, and to be 1 − *p*_*s*+1,*u*_ when *i* = *j* and they are not in the detached state. When both are in the detached state, *T*_*ij*_^(*s*+1,*u*)^ is 1, and for all other combinations of *i* and *j*, we set 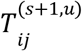 to 0.

#### N-terminal blocking rates

N-terminal blocking behavior is also defined by Bernoulli random variables. There are two of these rates, and we start by defining our factor-specific functions for the cyclic blocking behavior, which defines the rate at which a peptide moves into a blocked state instead of remaining unchanged.

We can only measure this rate from the unblocked states. Let *g*_*u*_ (*i*) = 1 in those states, and 0 for all other states. *h*_*u*_ (*i, j*) = 1 if *i* represents an unblocked state and *j* indicates the specifically corresponding blocked state. *h*_*u*_ (*i, j*) = 0 for all other combinations of *i* and *j*, though these should never occur under a correct implementation of this algorithm.

This again is a weighted maximum likelihood estimate of a Bernoulli random variable, and we can describe the action of ψ. 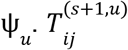 should be *T p*_*s*+1,*u*_ when *i* is an unblocked state and *j* is the corresponding blocked state, while it should be 1 − *p*_*s*+1,*u*_ when *i* = *j* and they are in an unblocked state. When both are in the same unblocked state, 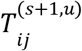 is 1, and for all other combinations of *i* and *j*, we set 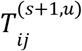 to 0.

For the matrix factor representing the initial N-terminal blocking, we analyze the data in exactly the same way, we just use a different parameter and matrix to track our results, in order to allow the initial blocking rate to be different from the cyclic one.

#### Fluorophore destruction before and during sequencing

An assumption of equal exposure to both chemical failure and photobleaching of the fluorophores means we should treat this as a binomial random variable. As with N-terminal blocking rates, we have one rate that identifies the behavior before sequencing starts (the missing fluorophore rate), and another for destruction during sequencing (the dye loss rate). These will again be mostly equivalent in their analysis, though tracked with different variables in order to enforce a separation. We show here the analysis for the dye loss rate. We also wish to emphasize that we track these separately, and with separate functions *g*_*u*_, *h*_*u*_, and ψ_*u*_, for each color of fluorophore.

We note that states with more fluorophores of the color being analyzed provide more evidence of the rate of fluorophore loss. With this in mind, *g*_*u*_ (*i*) is set to the number of fluorophores of the color of interest in state *i*. We then let *h*_*u*_ (*i, j*) = *g*_*u*_ (*i*) − *g*_*u*_ (*j*) if *j* ≤ *i*, and otherwise set it to 0 (for transitions which should never occur).

To construct *T*^(*s*+1,*u*)^, we let 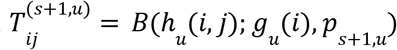 if *j* ≤ *i*, where *B* represents the parameterized probability mass function of the binomial distribution. This expands to:

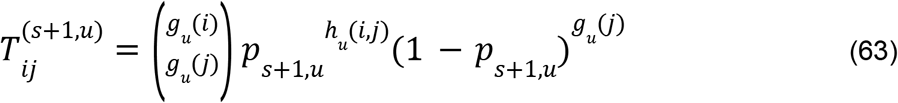

When *j* ≤ *i*. When *j* > *i*, we let 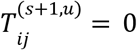.

There is a key difference in the analysis of the missing fluorophore rate. A bias is introduced because peptides with all fluorophores missing in every color will not be visible and are therefore absent from the data. A correction for this bias is discussed in (**Appendix A4**).

#### Edman degradation failure rate

Our factored transitions which manage Edman degradation are defined by a Bernoulli random variable, much like the factors managing the detachment rate or the two forms of the N-terminal blocking rates. While there is an additional complication from the probabilistic loss of a fluorophore in the case of a successful Edman degradation, we can safely ignore this until we need to construct *T*^(*s*+1,*u*)^. When determining *p*_*s*+1,*u*_, we need only concern ourselves with whether Edman degradation failed.

We need to omit states with a blocked N-terminus. For all states with a blocked N-terminus, *g*_*u*_ (*i*) = 0. For states with an unblocked N-terminus, *g*_*u*_ (*i*) = 1. We let *h*_*u*_ (*i, j*) = 0 if either or both of *i* or *j* represent a state with a blocked N-terminus. We also set it to 0 for all invalid combinations of *i* and *j* that should never occur; this relationship is complicated and peptide dependent. Combinations of *i* and *j* are valid when *i* = *j*, or when removing the N-terminal amino acid from state *i* can result in state *j*. In the second case, state *j* represents either the same combination of fluorophore counts of different colors but with one less amino acid, *or* it represents that combination less one amino acid *and* minus one fluorophore, *of the specific color* of fluorophore which may or may not be present on the amino acid being removed. This is invalid when the N-terminal amino acid cannot be labeled and in that case should be zero.

In any case, when *i* = *j*, we let *h*_*u*_ (*i, j*) = 0, as this is an Edman success. For all other *valid* combinations of *i* and *j*, we let *h*_*u*_ (*i, j*) = 1. This will give a weighted maximum likelihood estimate of the probability of an Edman failure event.

To define *T*^(*s*+1,*u*)^, we let 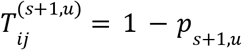 when *i* = *j*. For *i* ≠ *j* (assuming a valid transition), if state *i* does not have a possibility of a fluorophore on its N-terminal amino acid, then 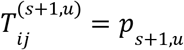. If a fluorophore is possible on its N-terminal amino acid, the computation is a bit more involved.

In [16], we derived a formula for the probability of fluorophore removal with Edman degradation, which we reiterate here under a slightly different symbolic representation. Let λ(*i*) represent the number of labelable amino acids which can take a fluorophore of the same color that the

N-terminal amino acid may have when in state *i*. Let *G*(*i*) represent the number of fluorophores of the same color the N-terminal amino acid may have, when in state *i*. We note that this second function is similar in form to the variant of *g* _*u*_ (*i*) described under “Fluorophore destruction before and during sequencing.” Now, in the case where *i* ≠ *j* (for a valid transition) and state *i does* have a possibility of an N-terminal amino acid, and *j*, in relation to *i*, represents an amino acid removal, we let:

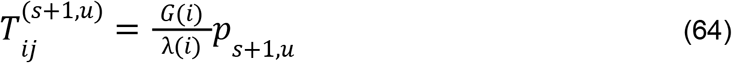

Then, for the case where *i* ≠ *j* (for a valid transition) and state *i* has a possibility of an N-terminal amino acid, and *j*, in relation to *i*, represents no amino acid removal, we then let:

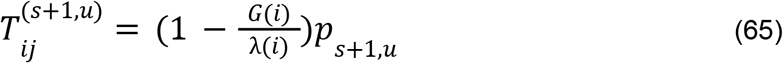

Finally, for invalid transitions, we let 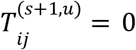.

### Additional discussion

We note that, with the exception of Edman degradation, all of the functions *g* _*u*_ (*i*) and *h*_*u*_ (*i, j*) are trivial to compute when the cumulative forward and backward probability results are indexed by a higher-order tensor, as described in [16]. Then, these functions become a simple extraction of an index of that higher-order tensor.

#### A4 -Bias correction for missing fluorophores

When a peptide is missing all its fluorophores it is not visible, and does not get sequenced. Correcting for this issue involves considerable additional complexity. We start by noting that this introduces a statistical dependency between missing fluorophore rates of different colors of fluorophores. This is because a count of zero fluorophores of one color only eliminates the peptide from the dataset if there are also zero fluorophores of every other color. We find it more clear to first define τ^(*s*+1,*u*)^. As in the case for the dye loss rate, let *g* _*u*_ (*i*) represent the number of fluorophores of the color of interest in state *i*, and let *h* _*u*_ (*i, j*) = *g* _*u*_ (*i*) − *g* _*u*_ (*j*) if *j* ≤ *i*, and otherwise set it to zero.

Now however, instead of defining τ^(*s*+1,*u*)^ with a parameterized binomial distribution as we did before for *T*^(*s*+1,*u*)^, we use a modified binomial distribution where the entries represent the probability of the transition *conditioned on* the probability of the peptide being observable. Letting Ĉ represent the set of all pre-sub-transition indices corresponding to missing fluorophore rates of different fluorophore colors, this equation is given by:

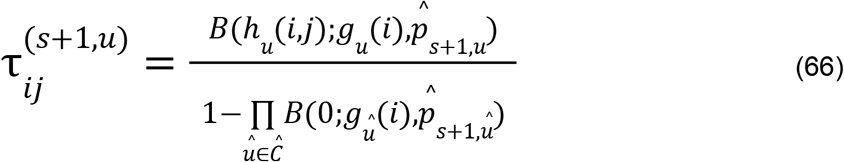

We of course require, as before, *j* ≤ *i*, and otherwise set 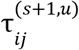 to zero. Additionally, we note that 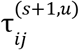 should be zero for the state *i* such that *g*_û_ (*i*) = 0 when û ∈ Ĉ (the state with no fluorophores of any color).

With other forms of error, we could point to existing and well understood formulas for their maximum likelihood estimates, as they can be viewed either as binomial or Bernoulli random variables. That will not work in this case, and we must explicitly derive this result. We consider the colors of fluorophore together, due to their statistical dependencies. Then we seek to maximize the likelihood given by:

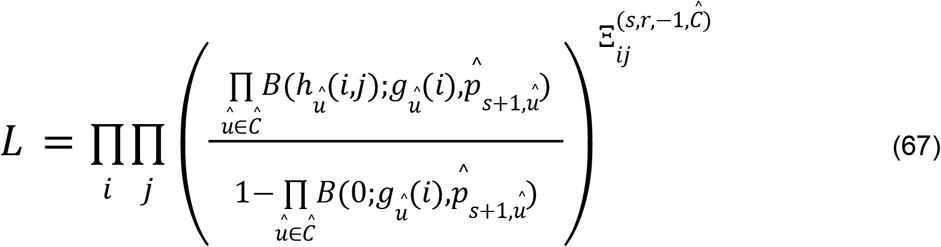

Assuming *i* and *j* vary over only their valid ranges, and where we let 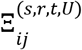 represent the probability of a transition from state *i* to state *j* given *Y*_1:*T*_ = *y*_1:*T*_ and θ^(*s*)^ (for iteration *s*, sequence *r*, time step *t*) when considering only the subset of sub-transitions contained in *U*, the set of all missing fluorophore related sub-sequences. In mathematical notation:

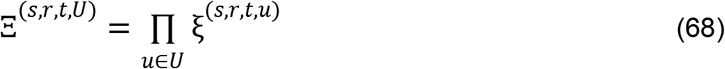

We now note a useful simplification to our formula. We previously defined our initial conditions for *f*^(*s*,−1)^ so that it would be 1 for the perfectly labeled and non-blocked state, and zero everywhere else. The only pre-sub-transition other than the missing fluorophore rate is the transition for initial N-blocking. The status of the N-terminus is irrelevant to the missing fluorophore rate, thus we consider the blocked and unblocked states together. We also ignore all states with missing fluorophores as irrelevant. We name the remaining state *I*. Then we can reduce our likelihood equation to:

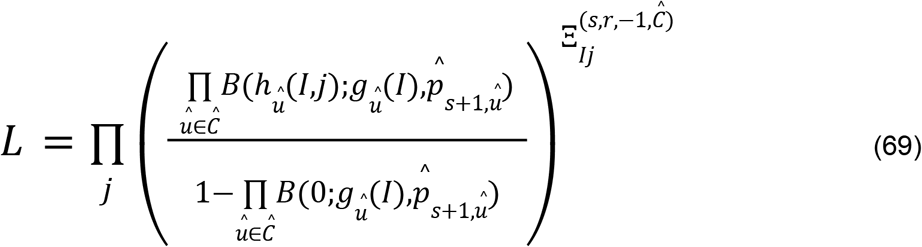

Expanding our equation for the likelihood we get:

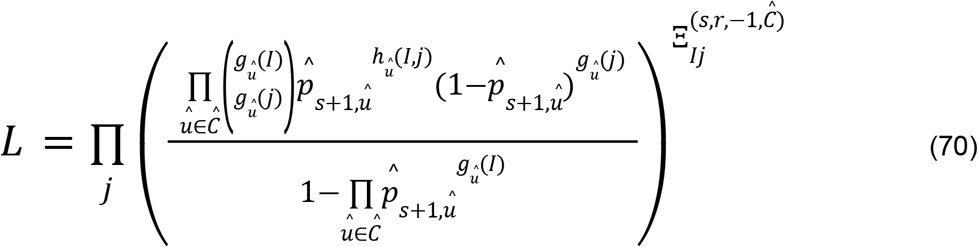

We now take the logarithm, which makes the equation easier to work with while preserving order:

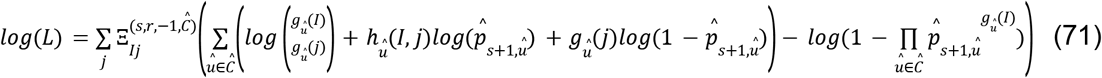

Noting that this equation is symmetric with respect to û, we need only solve for one result to maximize for every 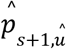. We then take the derivative with respect to a choice of 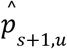 and set it to zero to search for extrema.

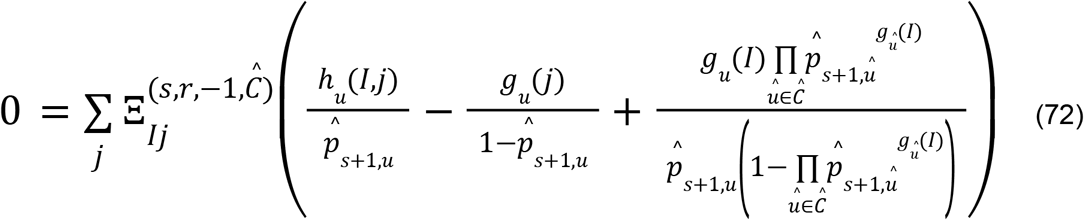

This reduces to:

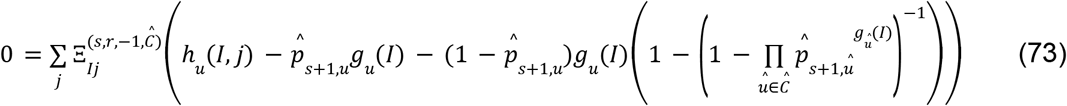

And reduces again to:

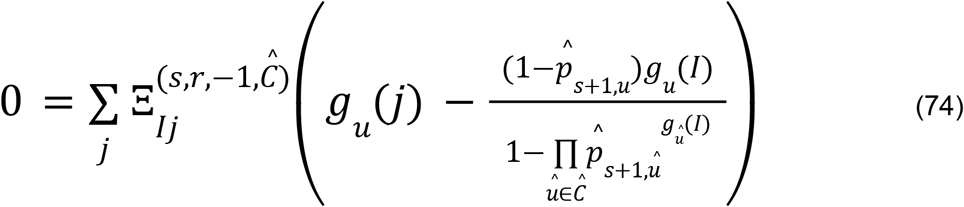

If we rearrange we find:

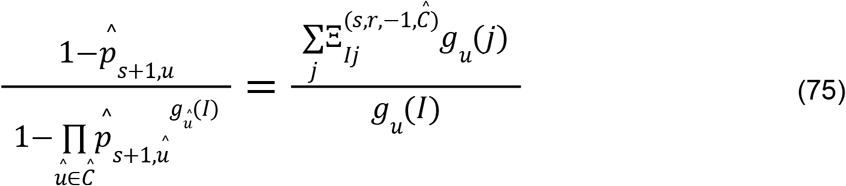

This is a multivariate generalization of the solution to a geometric sequence, and is therefore a generalization of the associated inverse problem. This also constitutes root-finding of a polynomial of arbitrary order, and thus is unlikely to have a closed form solution. If we assume large *g*_û_ (*I*) or small 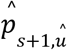 for at least one color of fluorophore, then this can be approximated by:

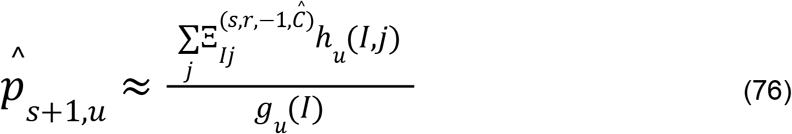

This is the maximum likelihood estimate for an ordinary binomial random variable. We use this to demonstrate that the extremum we’ve found is a maximum, in place of applying the second derivative test as would ordinarily be done.

We need a way to approximately solve this when *g*_û_ (*I*) is small or 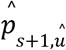 is large. We suggest an iterative method, where we plug the left hand side result into the right hand side on each iteration. We iterate over *z*, writing our new equation as:

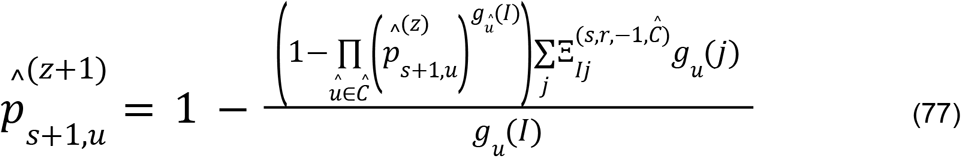

We need to prove that this iteration converges. The multivariate nature of this problem requires us to consider this in a multivariate vector space of dimension |Ĉ|. We also set some requirements. Firstly, this equation is clearly nonsense unless:

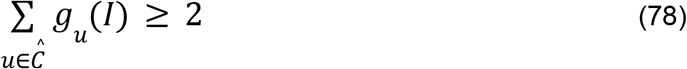

We also require a constraint on the solution:

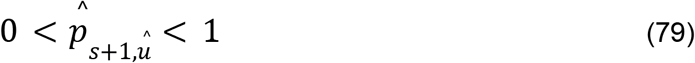

For convenience, we now introduce the following variables:

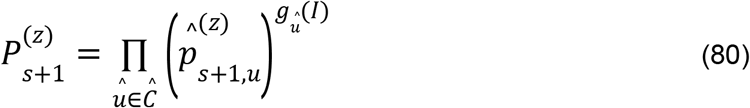

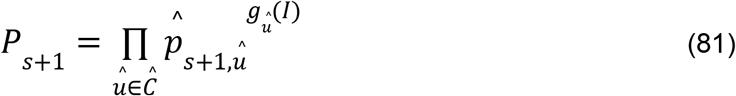

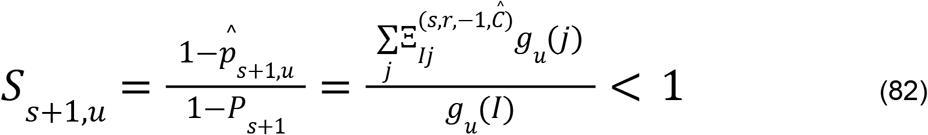

This gives us the equation:

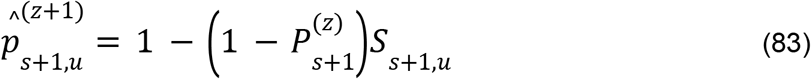

Equivalently:

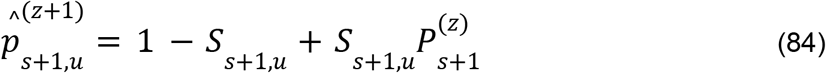

If we have a vector of 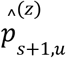 such that 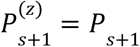 then clearly from this equation we will get the correct result in the next iteration, and 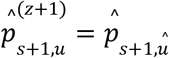. More generally, due to the linear relationship of these terms, and noting that *S*_*s*+1,*u*_ < 1, if 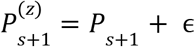 for |ϵ| ≪ 1, then there exists |δ| < |ϵ| such that 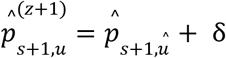. Therefore, to prove convergence of every 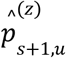 to 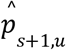 for all *u* ∈ Ĉ it is sufficient to prove convergence of 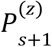 to *P*_*s*+1_.

There is a recurrence relation of 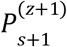 in terms of 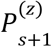, which is:

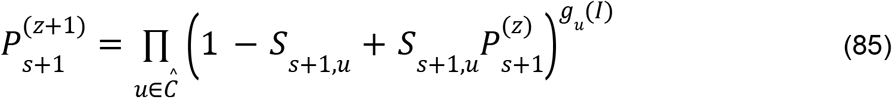

The right hand side is a polynomial expression of order 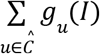 with all positive coefficients. It follows that its derivatives of every order up to and including 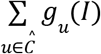 are strictly positive. In particular, our requirement that 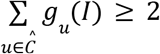 implies that the polynomial is strictly positive and has strictly positive first and second derivatives for 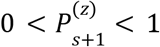. The strictly positive second derivative in this range guarantees that no more than two fixed points are possible in the given range. There is by definition a fixed point solution 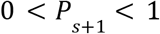, which is the solution we want our iteration to converge towards. There is also another fixed point at 1.

Noting that for 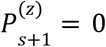 we get 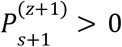, it follows that 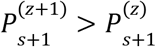 for 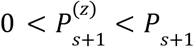. A double root at *P* _*s*+1_ is incompatible with the fixed point at 1. Therefore 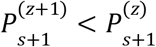 for 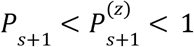. Using the strictly positive first derivative, we also find that 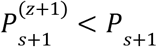 for 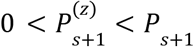 and we find 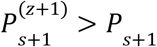 for 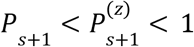. These results prove:

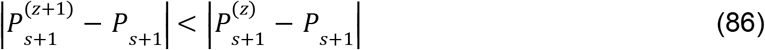

Therefore our iteration eventually converges towards the desired result for any 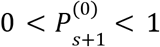.

We then fold this iteration into the outer iteration used to improve all estimates in the Baum-Welch algorithm, which gives the equation:

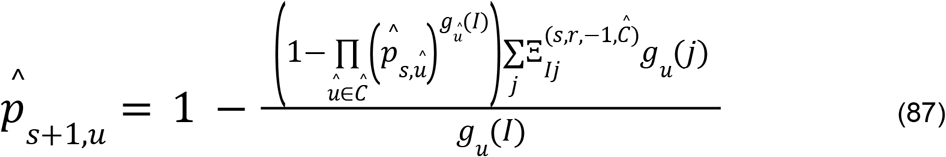

This is equivalent to imputing missing data using the parameter estimates in the previous iteration, as described in the main text. We also note that it is tempting to apply a root finding method from an open source library, which would likely converge faster in practice. However, our method allows us to accommodate cross-parameter effects by default, which is a desirable property. There is of course a valid concern that cross-parameter effects could in some way invalidate this proof, but we have found in practice that this does not appear to be the case.

